# Transcriptional and translational dynamics underlying heat shock response in the thermophilic Crenarchaeon *Sulfolobus acidocaldarius*

**DOI:** 10.1101/2022.12.17.520879

**Authors:** Rani Baes, Felix Grünberger, Sébastien Pyr dit Ruys, Mohea Couturier, Sarah De Keulenaer, Sonja Skevin, Filip Van Nieuwerburgh, Didier Vertommen, Dina Grohmann, Sébastien Ferreira-Cerca, Eveline Peeters

## Abstract

High-temperature stress is critical for all organisms and induces a profound cellular response. For Crenarchaeota, little information is available on how heat shock affects cellular processes and on how this response is regulated. In this work, we set out to study heat shock response in the thermoacidophilic model crenarchaeon *Sulfolobus acidocaldarius*, which thrives in volcanic hot springs and has an optimal growth temperature of 75°C. Pulse-labeling experiments demonstrated that a temperature shift to 86°C induces a drastic reduction of the transcriptional and translational activity, but that RNA and protein neosynthesis still occurs. By combining RNA sequencing and TMT-labeled mass spectrometry, an integrated mapping of the transcriptome and proteome was performed. This revealed that heat shock causes an immediate change in the gene expression profile, with RNA levels of half of the genes being affected, followed by the more subtle reprogramming of the protein landscape. A limited correlation was observed in differential expression on the RNA and protein level, suggesting that there is a prevalence of post-transcriptional and post-translational regulation upon heat shock. Furthermore, based on the finding that promoter regions of heat shock regulon genes lack a conserved DNA-binding motif, we propose that heat-shock responsive transcription regulation is likely not to be accomplished by a classical transcription factor. Instead, in contrast to histone-harboring Euryarchaeota that have heat-shock transcription factors, it is hypothesized that Sulfolobales and other histone-lacking thermophilic archaea employ an evolutionary ancient mechanism relying on temperature-responsive changes in DNA organization and compaction, induced by the action of nucleoid-associated proteins.

## Introduction

Crenarchaeota belonging to the *Sulfolobales* are characterized by a thermoacidophilic lifestyle with an optimal growth temperature between 75 and 80°C, enabling them to thrive in their natural habitat, geothermal hot springs (1). Because of the dynamic nature of a hot spring environment, cells are continuously confronted with temperature fluctuations. It can thus be anticipated that response to temperature stress, especially to heat shock, which compromises cellular integrity, makes up an inherent and important part of the stress response physiology of *Sulfolobales*. A sudden increase in temperature above the already high optimal growth temperature could lead to detrimental cellular damage, such as protein denaturation and aggregation, but also to increased membrane permeability and damage of nucleic acids (2). To counteract these effects, a heat shock response is initiated, consisting of a combination of universally conserved and (cren-)archaeal-unique features (3).

In *Sulfolobales*, heat shock response leads to an altered lipid composition of the cytoplasmic membrane (4, 5) and by the action of heat shock proteins (HSPs), which are universal key players in maintaining protein homeostasis by functioning as molecular chaperones (2, 3). Like in other archaea, and in contrast to eukaryotes and bacteria, the HSP machinery of *Sulfolobales* consists of a limited set of components (3). In addition to the HSP60-type group II chaperonin complex, also named thermosome, the HSP repertoire consists mainly of small HSPs (sHSPs) that typically oligomerize into diverse higher-order oligomers, functioning by directly binding to substrate proteins thereby protecting them from aggregation (6, 7, 8). The thermosome forms a large complex that refolds denatured proteins in an ATP-dependent manner. *Sulfolobales* encode three different thermosome subunits (α, β, and γ), with each subunit being differentially expressed and post-translationally regulated in response to temperature changes, leading to thermosome complexes with different subunit compositions and substrate specificities (9, 10). In response to heat shock, the thermosome complex of *Saccharolobus solfataricus* shifts from an octameric composition with alternating α and β subunits at physiological temperature to an all-β nonameric composition *in vitro* (10).

Given the far-reaching physiological effects of heat shock, it can be anticipated that an extensive gene regulatory program is induced, which most likely is not limited to differential expression of HSP subunits. Indeed, transcriptomic analysis has revealed that about one-third of all transcripts in *S. solfataricus* display a differential abundance at 5 minutes after reaching a stable heat shock temperature (11). Also, vapBC-like toxin-antitoxin systems have been shown to be differentially expressed and to play a role in heat shock response (11, 12, 13). The extensive heat-shock responsive transcriptional regulation was shown to be a transient mechanism, with most transcript levels returning to their baseline levels after prolonged heat shock (11). Moreover, also on a protein level and at longer time scales, regulation could be observed. Recently, a quantitative proteomics study was performed in the related species *Sulfolobus islandicus*, comparing growth at physiological and heat shock temperature (75°C and 85°C, respectively), demonstrating altered levels of a wide range of proteins belonging to different functional categories (14).

Regulatory mechanisms that underly heat shock response are well-understood in eukaryotic and bacterial systems, in which regulation is mostly dependent on global transcriptional regulators, such as the eukaryotic HSF1 activator or alternative σ factors in bacteria (*e*.*g*. σ^32^ in *Escherichia coli*) (2). In archaea, much less is known about heat-shock responsive gene regulation. The only exception is that it has been shown that certain Euryarchaeota harbor a dedicated transcription factor that exerts regulation in response to heat shock, such as Phr in *Pyrococcus furiosus* (15, 16, 17) and HSR1 in *Archaeoglobus fulgidus* (18). These regulators have a similar regulon, including a sHSP-encoding gene and a gene encoding an AAA+ ATPase involved in heat shock response. However, their regulons do not include genes encoding key HSPs such as the thermosome subunits, suggesting that other regulatory mechanisms might be involved in establishing a complex regulatory network (3).

As opposed to Euryarchaeota, the regulatory mechanisms underlying heat-shock responsive transcriptomic and proteomic dynamics are elusive in Crenarchaeota and more specifically in *Sulfolobales*. Moreover, while previous studies have revealed large alterations in transcript and protein levels in response to heat shock (11, 14), it is unknown whether these changes are mainly resulting from transcriptional, post-transcriptional and/or post-translational regulation or also by proteolytic and/or RNA degradation effects. In this study, we shed light on heat shock responsive gene regulation in the model crenarchaeal species *Sulfolobus acidocaldarius* by performing an integrated mapping of the transcriptomic and proteomic landscape. To this end, we combined RNA and protein pulse-labeling, RNA sequencing and TMT-labeled mass spectrometry methodologies. While improving the understanding of how *de novo* transcription and translation are affected by heat shock, we also reveal which key cellular processes are subjected to regulation and at which information processing levels regulation mainly takes place.

## Materials and Methods

### Strains, growth conditions and sample preparations

Starting from *S. acidocaldarius* strain SK-1 (19), genomic epitope tagging was performed to generate a strain expressing a FLAG-tagged thermosome α subunit, a 6xHis-tagged thermosome β subunit and a human influenza hemagglutinin (HA)-tagged thermosome γ subunit (SK-1*x*Thα-FLAG+Thβ-6xHis+Thγ-HA) (**Supplementary Methods**). All *S. acidocaldarius* strains used in this work (**Supplementary Table S1**) were cultured in Brock basal salts medium (1) supplemented with 0.2 % sucrose, 0.1 % NZ-amine and 20 μg ml^-1^ uracil. For RNA pulse-labeling experiments, the same medium was used but supplemented with 5 μg ml^-1^ uracil. For protein pulse-labeling experiments, Brock basal salts medium was supplemented with 0.2% sucrose, 10 μg ml^-1^ uracil and 0.73 g l^-1^ yeast methionine/uracil drop-out medium (CSM-Met-Ura, MP Biomedicals, USA, Solon) and lacked NZ-amine. All culture media were acidified to pH 3.0-3.5 with H_2_SO_4_. Cultures were grown at 75°C while shaking, unless stated otherwise, and growth was followed by measuring the optical density at 600nm (OD_600nm_). All primers used in this work are listed in **Supplementary Table S2** and an overview of plasmids is provided in **Supplementary Table S3**.

The experimental heat shock set-up for the omic-experiments was adapted from (20) (**Supplementary Figure S1**). Briefly, four biological replicates of *S. acidocaldarius* MW001 cultures were grown until mid-exponential phase (OD_600nm_ 0.45) after which they were transferred to a 6-well plate in a 75°C-preheated shaking heating block. The temperature of the heating block was increased to trigger a heat shock at 86°C and samples were collected *pre* and at different time points *post* heat shock. Each sample was split for transcriptomic and proteomic analysis. Samples for transcriptomic analysis were stabilized with an equal volume of RNAprotect® Bacteria Reagent (Qiagen, USA, Maryland) before centrifugation of all samples (10 minutes at 6,574 x *g* and 4°C). Samples for proteomic analysis were subsequently washed with 0.9 % NaCl. The SK-1*x*Thα-FLAG+Thβ-6xHis+Thγ-HA strain was included in the experimental set-up to validate the induction of a heat shock response. Detailed information about the samples is presented in **Supplementary Dataset S1**. Cellular viability was confirmed in heat-shock conditions by performing spot tests and heat shock induction was assessed by western blotting (**Supplementary Methods**).

### RNA sequencing and data analysis

RNA sequencing (RNA-seq) was performed by subjecting stabilized cell pellets to RNA extraction, followed by sample processing and high-throughput sequencing using an Illumina platform (**Supplementary Methods**). Data were processed by performing a quality control, mapping and differential expression analysis by EdgeR (21) (**Supplementary Methods, Supplementary Dataset S2** and **S3**).

### TMT-labeled Liquid-Chromatography-Tandem-Mass-Spectrometry and data analysis

Cell pellets were subjected to protein extraction, followed by further processing, Tandem Mass Tag (TMT) labeling and a Liquid-Chromatography-Tandem-Mass-Spectrometry (LC-MS/MS) analysis (**Supplementary Methods**). The resulting MS/MS data were further processed and differential expression analyzed by DEqMS (22) as described (**Supplementary Methods, Supplementary Dataset S3**).

### arCOG analysis, correlation analysis of RNA-seq and MS data and promoter analysis

Archaeal Clusters of Orthologous Genes (arCOGs) classification was retrieved from (23) followed by manual revision. Gene set enrichment analysis of arCOGs was performed using the goseq package in R, which accounts for gene length bias (24). Next, p-values for over- and under-representation of arCOG terms in the differentially expressed genes were calculated separately for up- and downregulated genes based on RNA-seq and MS data, respectively, and were considered as significantly enriched / not enriched below a cutoff of 0.05.

Pearson correlation coefficients were calculated from RNA-seq and MS log_2_ transformed count data and were used to analyze the association between the two methods.

Promoter motif analysis was performed by considering previously determined positions of primary transcription start sites (TSSs) (25). Sequences were extracted in a strand-specific way from - 50 to +1 nt from each TSS and plotted in R using the ggseqlogo package (26).

### Pulse-labeling of neosynthesized RNA

RNA pulse-labeling experiments were performed for *S. acidocaldarius* SK-1*x*Thα-FLAG+Thβ-6xHis+Thγ-HA cells cultivated to mid-exponential phase (OD_600nm_ 0.4) in low uracil medium, as described above. At different times before and after a heat shock from 75°C to 86°C, the culture was pulsed by adding either 4-thiouracil (4TU, 135 μM final concentration) or uracil (mock-control, 45 + 135 μM) (**Supplementary Figure S2**). After 30 minutes, 7-mL culture samples were centrifuged for 8 minutes at 4,000 x *g* and 4°C and pellets frozen at -20°C.

Total RNA was extracted from the cell pellets using the hot-phenol procedure and followed by biotinylation with MTSEA-Biotin-XX (Biotium, USA, Fremont) as previously described (27, 28). Northern blotting was performed by subjecting 5.5 μg biotinylated RNA to a denaturing agarose RNA gel electrophoresis at 20 V and transferring it to a positively charged nylon membrane by capillary transfer. Subsequently, biotinylated RNA was detected by DyLight™800-conjugated Streptavidin (Invitrogen, USA, Waltham) using the Odyssey CLx platform (LI-COR, USA, Lincoln) (27, 28). Streptavidin was stripped from the blot by pouring a boiling 1 % sodium dodecyl sulfate (SDS) solution twice over the membrane. The blot was prehybridized with 50 % formamide, 5x saline-sodium citrate (SSC, 750 mM NaCl, 75 mM sodium citrate, pH 7), 0.5 % SDS, 5X Denhardt’s Solution (1 mg ml^-1^ Ficoll 400, 1 mg ml^-1^ polyvinylpyrrolidone, 1 mg ml^-1^ bovine serum albumin, fraction V) for 3 hours at 30°C. For the detection of bulk steady-state rRNA, 10 pmol of fluorophore-coupled probes were added that target the 5’end region of mature 16S or 23S rRNA (DY682-CTTATCCCTACCCCGATAGCGG or DY782-CGAGCATTTCGCTGCTTGCCG, respectively), followed by overnight incubation at 30°C. Blots were first washed with 2X SSC during 15 minutes, then with 1X SSC during 15 minutes at 37°C and rRNA was visualized using the Odyssey CLx platform (LI-COR, USA, Lincoln). Fluorescence signals were quantified with ImageJ 1.53e (29).

### Pulse-labeling of neosynthesized protein

A Bioorthogonal Noncanonical Amino Acid Tagging (BONCAT) approach was used for protein pulse-labeling experiments. *S. acidocaldarius* SK-1*x*Thα-FLAG+Thβ-6xHis+Thγ-HA cells were cultivated to mid-exponential phase (OD_600nm_ 0.33) in absence of methionine, as described above. Next, heat shock and pulse-labeling was performed as described for RNA pulse-labeling, except that 1 mM L-azidohomoalanine (L-AHA) and 1 mM L-methionine were used as pulsing agent and mock-control, respectively (**Supplementary Figure S2**).

The BONCAT protocol employed in this study was slightly adapted from the one described for *Haloferax volcanii* (30). Briefly, proteins were extracted from the samples by boiling, reduced and 133 μg total protein per sample was alkylated in fresh PBS buffer with 0.2 M 2-chloroacetamide. L-AHA-containing proteins were labeled with Cy7 by strain-promoted azide-alkyne click-chemistry employing 1 μM dibenzylcyclooctyne-Cy7 (DBCO-Cy7) (Click Chemistry Tools, USA, Scottsdale). After methanol-chloroform extraction, protein pellets were resuspended in 40 μL of 2x HU-buffer (100 mM Tris-HCl pH 6.8, 0.5 mM EDTA pH 8, 4 M Urea, 2.5% SDS) supplemented with 200 mM dithiothreitol (DTT). Protein aliquots of 5 μL were loaded on SDS-PAGE and separated at 180 V. Fluorescence signals of L-AHA-incorporated proteins were detected in-gel using Odyssey CLx platform (LI-COR, USA, Lincoln). Bulk, steady-state levels of protein were detected by Coomassie-staining of the gel and scanning employing the Gel Doc XR+ System (Bio-Rad, USA, Hercules). Signals were quantified with ImageJ 1.53e (29).

### Data availability

RNA-seq data were deposited in the European Nucleotide Archive at EMBL-EBI under accession number PRJEB57647. MS data were deposited to the ProteomeXchange Consortium via the PRIDE (31) partner repository with the dataset identifier PXD038744.

Convenient, easily accessible lists of genes of pathways with their corresponding expression levels are available in **Supplementary Dataset S3**. Code for all bioinformatic analyses is available via Github (github.com/felixgrunberger/HSR_Saci).

## Results

### Global transcriptomic and proteomic changes in response to heat shock

When studying heat shock response, it is imperative to subject cells to a temperature change that elicits a stress response without affecting cellular integrity. At mid-exponential growth phase, a rapid temperature shift from 75 to 86°C (**Supplementary Figure S1**) was shown to induce a regulatory response in *S. acidocaldarius*, as demonstrated by the upregulation of the thermosome α and β subunits (**Supplementary Figure S3a**), without impairing cellular viability (**Supplementary Figure S3b**). With the aim of simulating how temperature gradients are sensed in the natural environment, we selected three time points after shifting cells to 86°C (**Figure 1a**). These represent conditions in which cells are still experiencing a temperature increase (15 minutes), in which a steady-state has just been reached (30 minutes) and in which cells are subjected to a persistent heat-shock condition (60 minutes).

**Figure 1.**
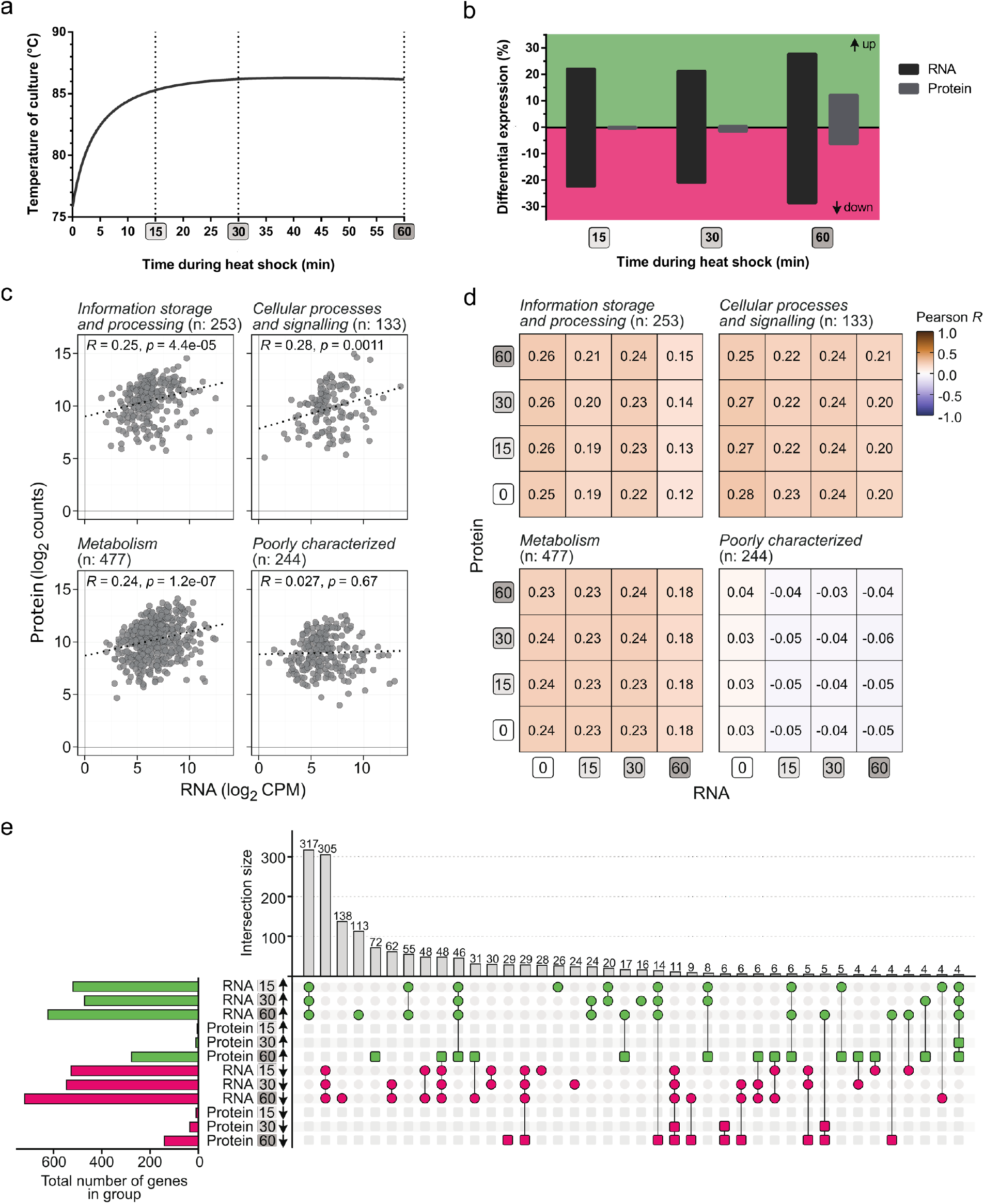
Global transcriptomic and proteomic changes in response to heat shock. **a**. Temperature profile of the culture upon temperature shift from 75°C to 86°C, according to the heat shock set-up described in (20). **b**. Differential expression analysis depicting the percentage of differentially expressed/produced genes and proteins at different *post* heat-shock treatment time points, compared to levels at 75°C. **c**. Correlation plot of the transcript and protein abundance at 75°C, subdivided according to the functional category of the respective gene. The Pearson correlation coefficient R is given for each category. **d**. Correlation matrix plotting the Pearson correlation R for transcript and protein abundances at 75°C and the heat shock time points (and the combinations thereof). **e**. Plot visualizing the overlap (intersection) between the regulation groups (increased or decreased abundance at 15, 30 and 60 minutes after heat shock treatment at the RNA or protein level). The size of the regulation groups corresponds to the number of differentially expressed genes/proteins in that group (related to panel b).

Upon subjecting cultures to transcriptomic and proteomic analyses, it was confirmed that massive, global changes occur at a transcriptomic level in response to a temperature increase, while this response is much less pronounced at a proteomic level (**Figure 1b**). Indeed, immediately after heat shock, 44.5% of all coding genes displayed a differential RNA abundance (1,047 of 2,351 genes), decreasing to 42.2% (991 genes) upon reaching a steady state and increasing again to 56.4% at 60 minutes *post* heat-shock treatment (1,325 genes). The fraction of proteins displaying differential abundance evolved from 0.7% (15 of 2,267 proteins) at 15 minutes to 2.0% (46 proteins) at 30 minutes and 18.4% (418 proteins) at 60 minutes *post* heat-shock treatment (**Figure 1b**). While the number of genes with a higher and lower RNA level was approximately the same for the different time points, there was a bias towards a larger fraction of proteins with higher than lower abundance after 60 minutes (**Figure 1b**).

The strongly differing numbers of transcripts and proteins with differential abundance suggest that there might be a limited overlap between transcript- and protein-level effects, which was assessed by a gene-by-gene correlation analysis (**Figures 1c** and **d**). Indeed, a lack of correlation was confirmed by comparing the abundance of a particular transcript and its corresponding protein at 75°C and within the same heat shock time point, but also when comparing RNA levels at *e*.*g*. 15 or 30 minutes with protein levels at 60 minutes. A time-delayed reaction of the proteome in the same direction as the transcriptome does not seem to be a general pattern of the heat shock response in *S. acidocaldarius* (**Figure 1e**). These observations suggest a prevalence of post-transcriptional and post-translational regulation upon heat shock in addition to the extensive transcriptional regulation that was observed.

### Functional categories of differentially expressed genes

With the aim of exploring the functional context of genes and proteins that display a heat-shock responsive differential abundance, we performed a gene set enrichment analysis based on the arCOG classifications (23) (**Figure 2**). It is apparent that transcriptional heat-shock responsive regulation was not restricted to a limited set of functional categories, but that it affected diverse biological processes. For information storage and processing gene categories, such as transcription and translation, considerable transcriptional downregulation was observed at most time points (**Figure 2**). In addition, an overrepresentation of downregulated transcripts and proteins was found within categories such as cell cycle control, cell division, chromosome partitioning, cell motility and signal transduction mechanisms. In contrast, genes involved in post-translational modification and protein turnover are significantly overrepresented in upregulated transcripts and proteins. Also, several metabolic pathways were affected, including pathways in nucleotide and amino acid metabolism, energy production and carbohydrate metabolism (**Figure 2**).

**Figure 2.**
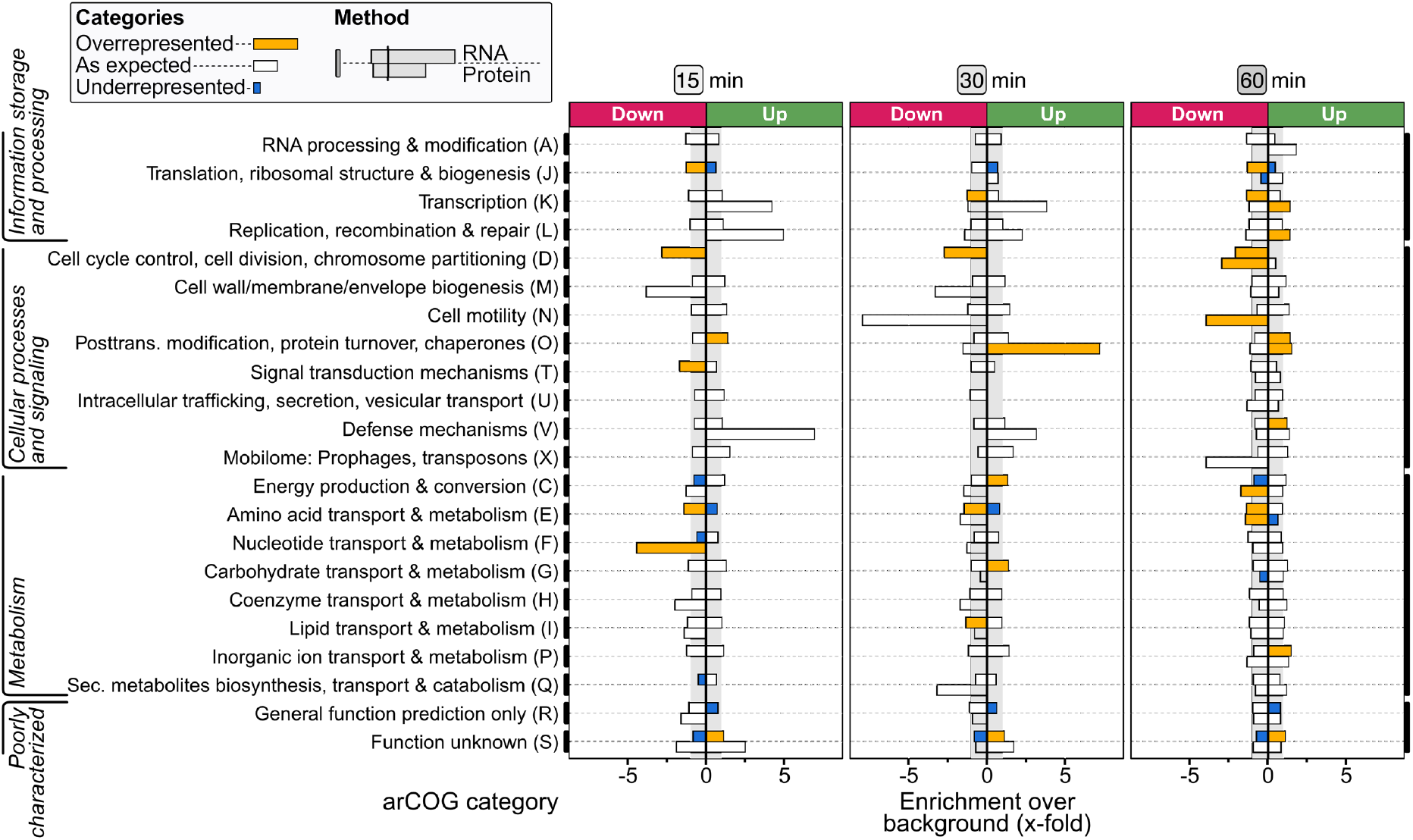
Gene set enrichment analysis of archaeal clusters of orthologous genes (arCOGs). Bars indicate the fold enrichment of increased (up) or decreased (down) abundance of genes (top bar) and proteins (bottom bar) for each arCOG category, based on the total number of differentially expressed genes/proteins at that time point and by incorporating the number of genes/proteins belonging to that particular arCOG category. Bars are color*-*coded according to statistically significant over- or underrepresentation (p < 0.05). Note that one should be careful with interpretation of the protein data at 15 and 30 minutes, given the low number of differentially produced proteins at these time points (see **Figure 1b**).

### Response of the heat shock protein machinery

The most apparent upregulation was observed for the arCOG category O, comprising genes involved in post-translational modification, protein turnover and chaperones, with a significant overrepresentation of upregulation on both the RNA and protein level at 60 minutes *post* heat shock (**Figure 2**). This category includes the HSP machinery and proteasomal degradation system as key elements in the preservation of structural integrity of proteins (**Figure 3, Supplementary Results**).

**Figure 3.**
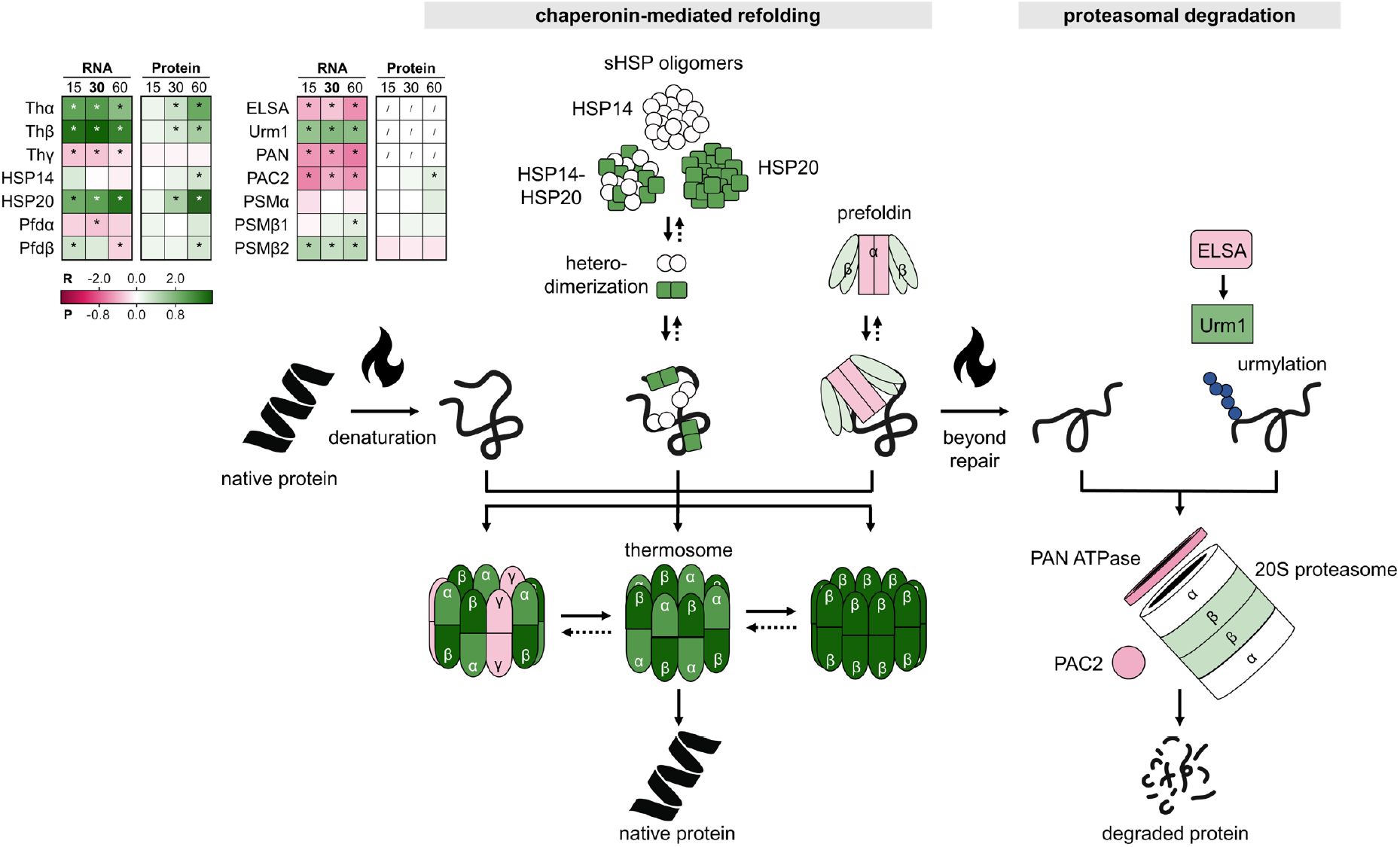
Heat-shock responsive differential abundance of the chaperonins or heat shock proteins (HSPs) and the proteasome. The inset table shows differential expression at the RNA (R) and protein (P) level at all time points (15, 30, 60 minutes), color-coded according to the log_2_FC value in the gradient below. * = significant (FDR/adj.p-value < 0.05). / = not covered. The scheme on the right is colored according to differential expression at the RNA level at 30 minutes.

During exponential growth at physiological conditions at 75°C, all HSPs were expressed both at the RNA and protein level (**Supplementary Dataset S3**). Moreover, the three thermosome subunits α, β and γ were found to be amongst the four most abundant proteins in the cell (**Supplementary Figure S4**), underlining the importance of this chaperone complex in proteome homeostasis and cellular physiology even in absence of temperature stress. In response to heat shock, a substantial increased transcriptional gene expression was observed for the α and β subunits, followed by a slower increase in their protein levels, while the γ subunit was not heat-shock responsive (**Figure 3**). This finding is in agreement with the western blot analysis of the thermosome-tagged strain (**Supplementary Figure S3a**) and previous findings in *S. acidocaldarius* and *Saccharolobus shibatae* (9, 20, 31). In addition, the sHSPs, HSP14 and HSP20, were both upregulated, with HSP20 being the most highly upregulated protein after 60 minutes of heat shock in terms of differential expression (**Supplementary Dataset S3)**.

A mixed view emerges for the proteasome and associated proteins, with some of the components that were upregulated (*e*.*g*. a slight and considerable transcriptional increase was observed for the proteasome β subunit and Urm1, respectively) and others that were downregulated (*e*.*g*. a steady decrease was observed for transcripts encoding the proteasome-activating nucleotidase (PAN) AAA^+^ ATPase and the proteasome assembly chaperone (PAC2) (**Figure 3**). It is possible that urmylation modifications are increased upon heat shock, despite a lack of detecting Urm1 at the protein level. However, a lack of detecting a considerable upregulation of the essential proteasome components PAN

ATPase and PAC2 suggests that the effects of the high-temperature stress on the protein level can still be compensated by chaperones and that additional proteolytic support is not (yet) required.

More detailed descriptions of observations for these and other cellular processes (DNA topology, motility and biofilm formation, DNA replication, genome segregation and cell division, DNA repair and DNA import, post-transcriptional and post-translational modifications) are provided in **Supplementary Results** and **Supplementary Figures S5-S7**.

### Effects of heat shock on the functioning of the transcription process

The observation of a differential transcript or protein abundance does not necessarily result from changes in transcriptional or translational activity. However, it could also be the consequence of changes in RNA or protein stability, thereby affecting its degradation. To investigate this possibility, we analyzed the impact of heat shock treatment on the overall transcriptional and translational activity in *S. acidocaldarius* using a pulse-labeling approach.

For the analysis of transcriptional activity, the uracil analog 4TU (27, 28) allowed to follow the *de novo* synthesis of RNAs before and after heat shock (**Figure 4a** and **Supplementary Figure S8**). The major transcriptional activity of the cell comprises rRNA synthesis, and thus, as expected, 16S and 23S rRNA were the major species of neosynthesized RNA, both during growth at 75°C and after heat shock treatment. In addition, several larger RNA species were neosynthesized within a 30-minute time frame, which are hypothesized to be mRNA species with a high abundance and/or turnover rate, given that they are not hybridized by rRNA probes (**Supplementary Figure S8a**). Immediately after heat shock, a drastic decrease was observed for 4TU incorporation in rRNAs down to 20-40% of the initial levels at 75°C. The low level of neo-synthesized rRNAs was maintained for at least two hours after heat shock (**Figure 4a** and **Supplementary Figure S8b**), suggesting that heat shock causes the transcriptional activity to decline quickly.

**Figure 4.**
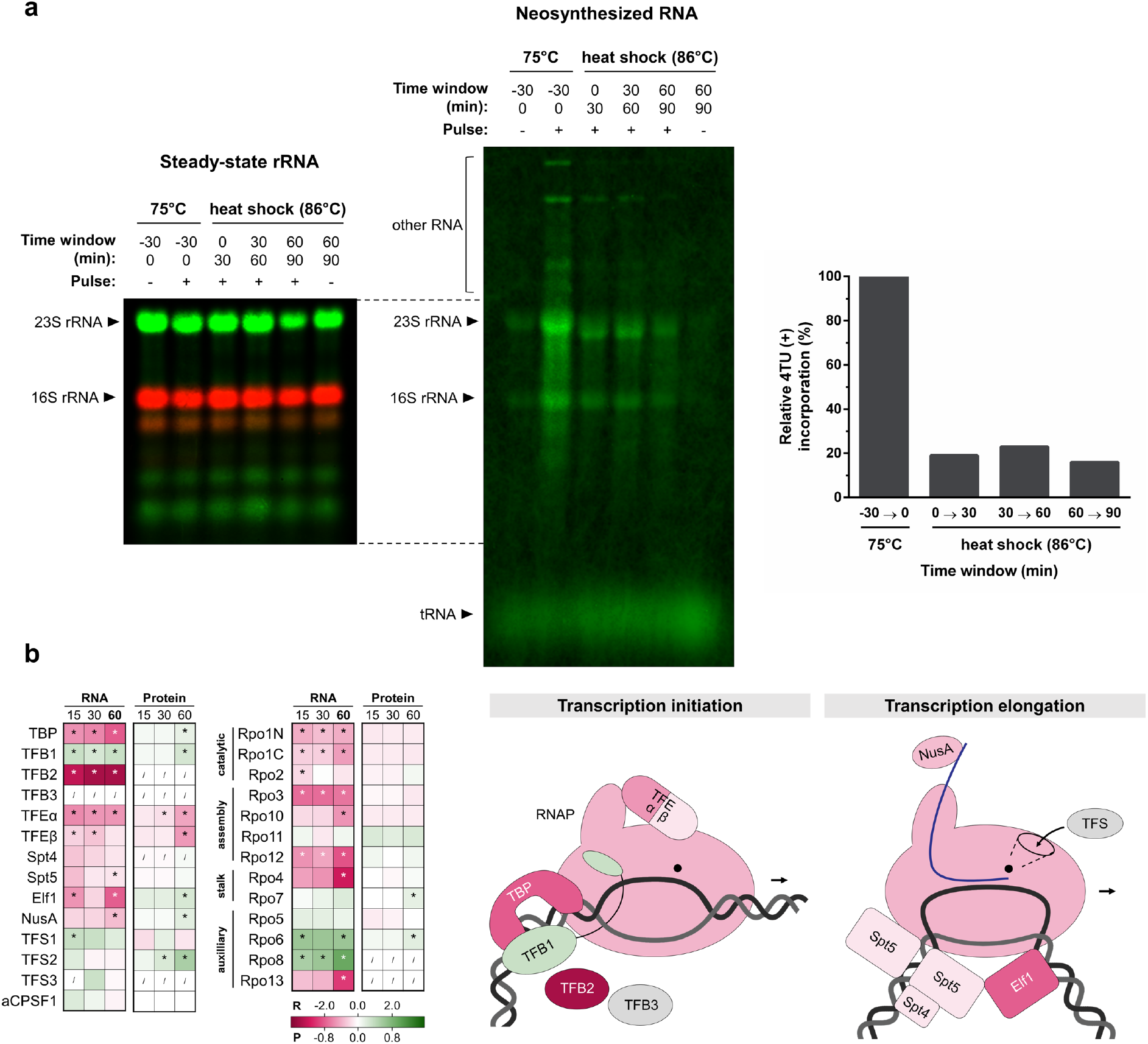
Impact of heat shock on transcription. **a** Pulse-labeling of neosynthesized RNA. SK-1 Thα-FLAG + Thβ-6xHis + Thγ-HA cultures were pulsed, both at 75°C and upon an 86°C heat shock for 30-minute time windows by addition of an excess of nucleotide analog 4TU (+) or uracil (-) as mock-control. 4TU was incorporated into neosynthesized RNA and 4TU-labeled RNA was biotinylated in the total pool of RNA. Total RNA was analyzed by Northern blotting. Bulk steady-state rRNA was detected by DY682- or DY782-coupled probes targeting the 5’end region of mature 16S or 23S rRNA, serving as a loading control (left panel). 4TU-labeled RNA was detected by DyLight800-conjugated streptavidin (central panel). Incorporation of 4TU into 16S or 23S rRNA was quantified by first subtracting the background-signal in the uracil-mock control and dividing this by the signal of the steady-state rRNA. Relative 4TU-incorporations upon heat shock were calculated relative to incorporation at 75°C and the average of 16S and 23S rRNA was determined as a ratio to express the relative transcriptional activity upon heat shock (right panel). **b** Differential expression of the transcription machinery upon heat shock. The inset table shows differential expression at the RNA (R) and protein (P) level at all time points (15, 30, 60 minutes), color-coded according to the log_2_FC value in the gradient below. * = significant (FDR/adj.p-value < 0.05). / = not covered. The graphical scheme on the right is colored according to differential expression at the RNA level at 60 minutes and is adapted from (33).

A decline in transcriptional activity is also supported by a downregulation for most of the individual components of the basal transcription machinery on the transcriptional level. However, this downregulation was not observed for all genes on the protein level: here, differential abundance and regulatory trends were less pronounced (**Figure 4b**). For example, immediately after heat shock, transcript levels were found to be decreased significantly for TATA binding protein (TBP), for the catalytic, assembly and stalk subunits of RNA polymerase (RNAP) and for the α and β subunits of basal transcription factor TFE (**Figure 4b**).

On the protein level, a downregulation was also observed, but only in a significant manner for TFEα and β, while most RNAP subunits were only slightly downregulated. TBP displayed an opposite effect, with a lower transcript and higher protein abundance. In contrast, TFB1, which is considered as the housekeeping TFB in *S. acidocaldarius*, was slightly upregulated on both the transcript and protein level, while the alternative TFB-type factor TFB2, of which the function remains enigmatic (33), displayed a strong transcriptional downregulation and was undetectable on the protein level. Taken together, transcription elongation factors exhibited a trend towards transcriptional downregulation upon heat shock with exception of TFS2, for which a considerable upregulation was observed.

### Effects of heat shock on the functioning of the translation process

Pulse-labeling experiments were also performed to map *de novo* translation by making use of the methionine analog L-AHA (30). Overall translational activity decreased after the temperature shift. However, this heat-shock responsive decline proceeded linearly and at a slower pace than the decline in transcriptional activity (**Figure 5a**). Contrary to this general trend, a small subset of abundant proteins was produced at higher levels upon heat shock. For example, this was the case for a protein represented by a noticeable band at 60 kDa, which could be hypothesized to represent the thermosome α and/or β subunit, as was also observed by western blotting in the thermosome-tagged strain (**Supplementary Figure S3a**).

**Figure 5.**
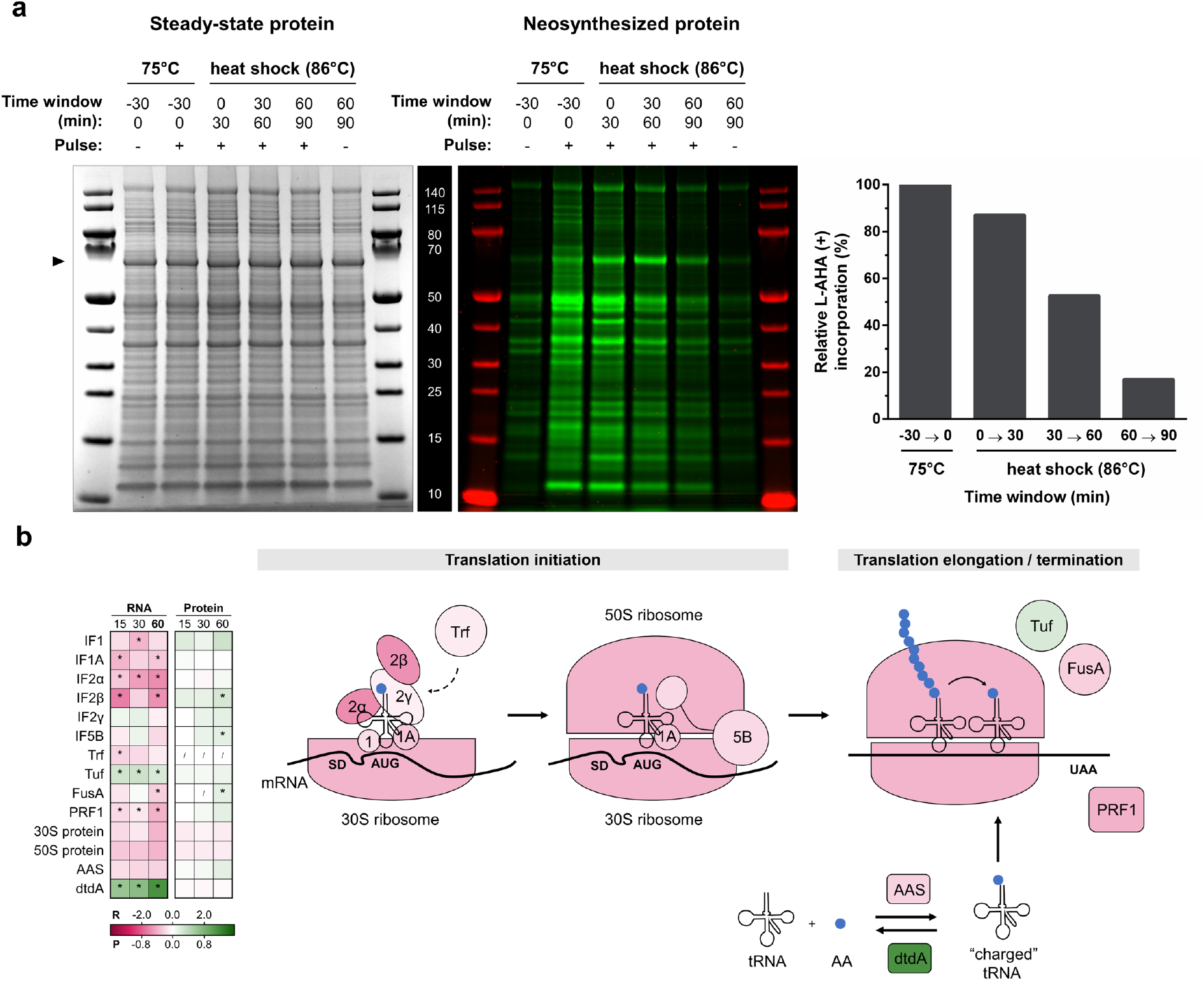
Impact of heat shock on translation and the translation machinery. **a** Pulse-labeling of neosynthesized protein. *S. acidocaldarius* SK-1 Thα-FLAG + Thβ-6xHis + Thγ-HA was pulsed at 75°C and upon heat shock for 30-minute time windows by addition of an excess of amino acid analog L-AHA (+) or methionine (-) as mock-control. Total proteins were extracted, reduced and alkylated. L-AHA-incorporated proteins in the total protein pool were labeled with Cy7 by strain-promoted azide-alkyne click-chemistry. Total proteins were separated by SDS-PAGE. Bulk, steady-state levels of protein were detected by Coomassie-staining (left panel), serving as a loading control. L-AHA-incorporated proteins were detected in-gel by Cy7-fluorescence detection (central panel) and quantified with respect to total protein (right panel). **b** Differential expression of the translation machinery upon heat shock. The inset table shows differential expression at the RNA (R) and protein (P) level at all time points (15, 30, 60 minutes), color coded according to the log_2_FC value in the gradient below. The median value of 30S, 50S proteins and L-aminoacyl-tRNA synthetases (AASs) is shown. * = significant (FDR/adj.p-value < 0.05). / = not covered. The scheme on the right is color-coded according to differential expression at the RNA level at 60 minutes and is adapted from (34)(35).

As observed for the basal transcription machinery, most components of the translation machinery were found to be transcriptionally downregulated upon heat shock, while this regulatory trend was not carried on at the protein level (**Figure 5b**). Indeed, most translation initiation factors were transcriptionally downregulated at the different time points, with IF2β and IF5B exhibiting the opposite effect -a significant upregulation-on the protein level at the 60-minute time point. With respect to translational elongation, Tuf displayed a slightly increased RNA abundance, while FusA displayed a slightly decreased RNA abundance and increased protein abundance. The translational termination factor PRF1 was transcriptionally downregulated. In addition, the RNA levels of most L-aminoacyl-tRNA synthetases (AASs), responsible for coupling of the correct amino acid to tRNAs, decreased (**Supplementary Dataset S3**). In contrast, D-aminoacyl-tRNA deacylase (dtdA), involved in recycling toxic D-aminoacyl-tRNA, was transcriptionally strongly induced during heat shock treatment (**Figure 5b**).

Considerable changes were observed in the expression of several components of tRNA and rRNA maturation pathways (**Supplementary Figure S9**). More specifically, CCA tRNA nucleotidyltransferase, which builds and repairs the 3’-terminal CCA sequence of tRNAs, and a 2’-5’ RNA ligase, both transcribed in the same operon, were among the most highly transcriptionally upregulated genes upon heat shock treatment. CCA tRNA nucleotidyltransferase is also strongly upregulated on a protein level at 30 and 60 minutes after heat shock. In addition, also RNase Z and RNA ligase RtcB are strongly upregulated at the RNA level (**Supplementary Figure S9**). Additionally, many enzymes involved in tRNA modification are differentially expressed after heat shock, albeit to a smaller extent (**Supplementary Dataset S3**). These observations indicate that although translational activity itself is slowed down upon heat shock, auxiliary processes that ensure translational fidelity and RNA processing are maintained or even upregulated.

### Search for putative heat-shock responsive cis-regulatory elements

In order to infer a putative DNA-binding recognition motif for a presumptive heat shock transcription factor, we screened promotor regions of transcripts with the most significant differential expression. Analysis in a position-unspecific manner did not yield any significantly enriched motifs in any of the regulation groups, which is furthermore supported by the plotted consensus motifs in these promoter regions (**Figure 6a**). Despite the absence of a transcription factor binding motif, it was apparent that strongly upregulated transcripts (especially at 15 and 30 minutes *post* heat shock) harbor more highly conserved BRE and PPE promoter elements (**Figure 6a** and **Supplementary Figure S10**). Given that this subset of genes did not have a significantly higher constitutive level of transcriptional expression in optimal growth conditions at 75°C (**Figure 6b**), a strong BRE and PPE promoter element seems to be an adaptation of highly induced heat-shock responsive genes and is linked to the regulation and not to basal transcription, possibly through the action of the TFB and/or TFE basal transcription factors.

**Figure 6.**
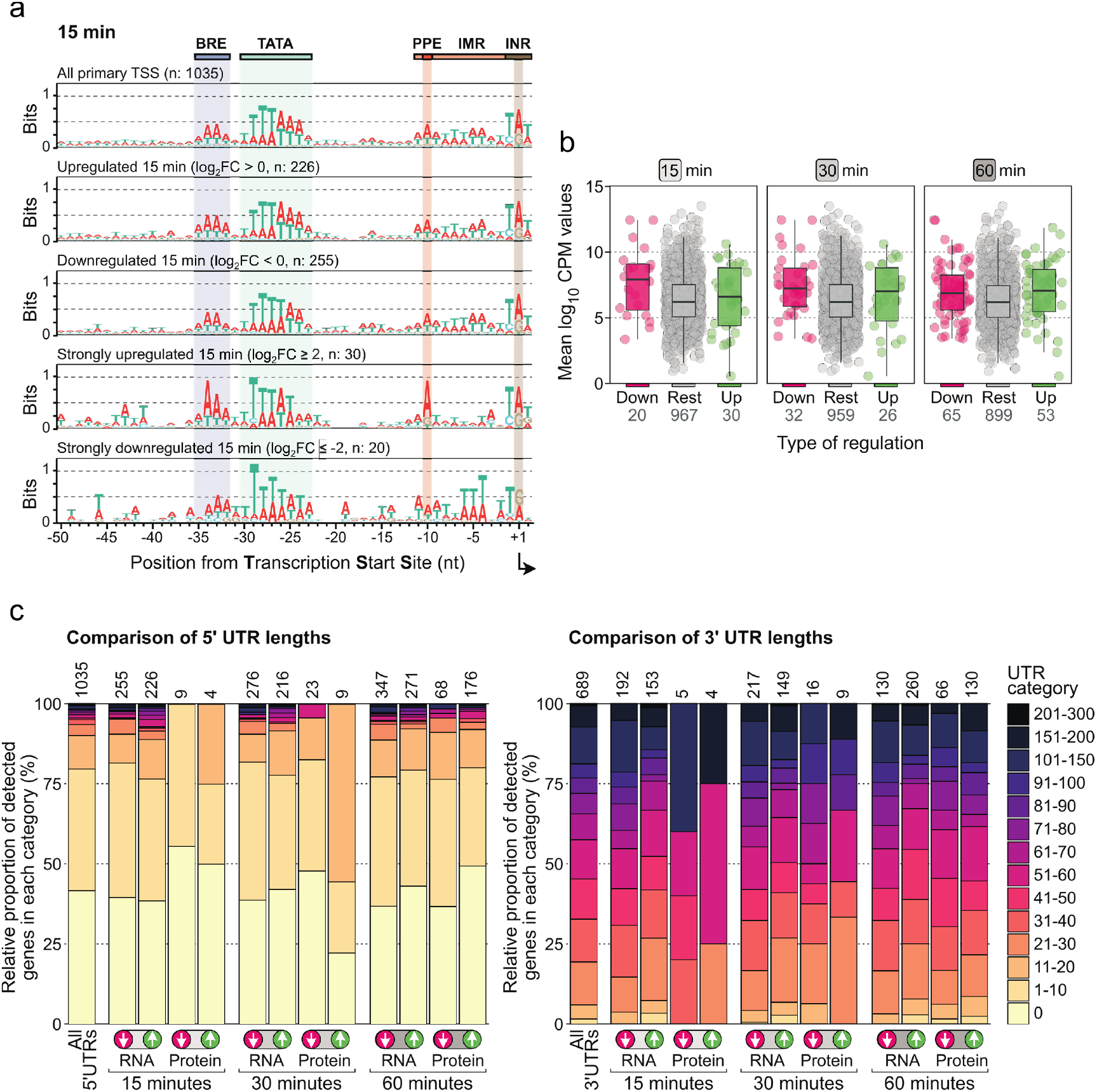
Search for putative heat-shock responsive *cis*-regulatory elements in promoter and UTR regions. **a**. Promoter motifs of all primary transcripts and subsets of regulation groups at 15 minutes after heat shock. PPE = proximal promoter element. IMR = Initially melted region. INR = Initiator element. **b**. The abundance of a given transcript at 75°C is plotted, subdivided according to the regulation groups at the different heat shock time points. Rest = not differentially expressed. High or low transcript levels at 75°C are not a prerequisite for strong up- or downregulation (log_2_FC ≥ 2 or ≤ -2) after heat shock. **c**. Bar plot showing the proportion of UTR-lengths (5’UTR on the left panel and 3’UTR on the right panel) for all genes in standard growth conditions (left bar) and for the differentially expressed transcripts/proteins at the different heat shock time points. Information on 5’UTR length was retrieved from (25) and 3’UTR length from (39).

In a next step, we assessed whether a correlation exists between 5’ and 3’ UTR lengths of mRNAs and their differential abundance on RNA or protein level (**Figure 6c**). Most transcripts in *S. acidocaldarius* are leaderless and lack a 5’-UTR (25, 35). However, it could be hypothesized to be a hot-spot region for temperature-responsive regulatory elements, as found in bacterial systems (*e*.*g*. RNA thermosensor element in the σ^32^ 5’-UTR (36, 37)). No correlation was found for 3’ UTRs on both the RNA and protein level and for 5’ UTRs on the RNA level (**Figure 6c**). On the protein level, however, proteins originating from transcripts with a 5’-UTR were overrepresented (higher protein abundance at 30 minutes after heat shock). It should be noted that this is based on a very limited set of genes: only 9 proteins displayed a higher abundance in these conditions, of which 5 (55.6%) with a 5’-UTR length between 10 and 20 nucleotides (nts), as opposed to 10.5% for all genes (**Figure 6c**). Notably, these 5 transcripts encode the thermosome α subunit (*Saci_0666*), thermosome β subunit (*Saci_1401*), HSP20 (*Saci_0922*), an Lrs14-like DNA binding protein (*Saci_1223*) and TFS2 (*Saci_1587*). Among these, several fulfill key functions in heat shock response.

### Effects of heat shock on chromatin organization

During the one-hour heat shock, we observed considerable changes in expression level of genes and proteins involved in chromatin organization (**Figure 7, Supplementary Results**). The macro-level chromosome organizer coalescin (ClsN) was slightly more abundantly expressed after heat shock, both on the transcriptional level (15-60 minutes) and the protein level (60 minutes) (**Figure 7a**). Focusing on the two Alba paralogs, we observed a late transcriptional downregulation of *alba-1* (60 minutes) although *alba-2* RNA levels were strongly decreased immediately upon heat shock (15-60 minutes), while remarkably, its protein level was increased (60 minutes). This suggests that there might not be an increase in Alba-1-Alba-2 heterodimers (40), but that Alba-2-regulated chromosome organization plays a role in heat shock response.

**Figure 7.**
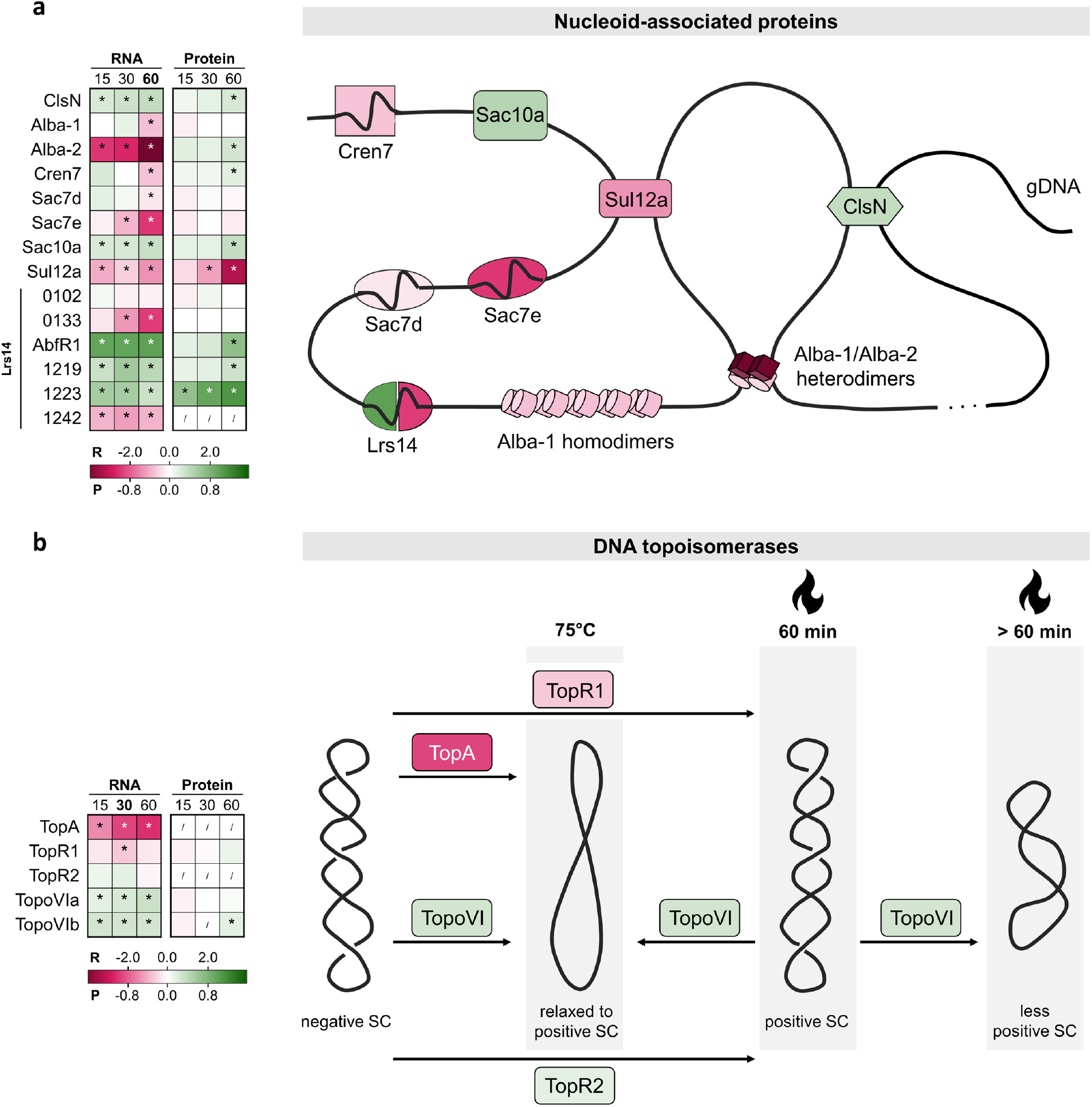
Heat-shock responsive differential abundance of genes/proteins involved in chromatin organization. The tables show the differential expression of the genes involved at the RNA (R) and protein (P) level at all time points after heat shock (15, 30, 60 minutes), color coded according to the log_2_FC value in the gradient below. * = significant (FDR/adj.p-value < 0.05). / = not covered. **a** Nucleoid-associated proteins. The figure shows the proteins in their cellular context, colored according to differential expression at the RNA level after 60 minutes. **b** DNA topoisomerases. The figure shows the proteins in their cellular context, colored according to differential expression at the RNA level after 30 minutes. Scheme inspired by (43,44, 46).

All small nucleoid-associated proteins (NAPs), except *sac10a*, displayed a decreased abundance at the RNA level upon heat shock, although this was not reflected on a protein level (**Figure 7a**). In contrast, most NAPs displayed a slight but significant increase in protein levels. Looking at the Lrs14 family of NAPs, which were previously shown to help controlling DNA topology after heat shock and stabilize DNA against thermodenaturation (40, 41, 42), it was observed that three out of six members of this family displayed an increase in transcript and protein levels (**Figure 7a**). Moreover, it is notable that *Saci_1223* is one of the most upregulated proteins at all time points and AbfR1 at 60 minutes (**Supplementary Dataset S3**).

Finally, also DNA topoisomerases displayed a differential transcriptional abundance in response to heat shock, although this was less pronounced on the protein level (**Figure 7b, Supplementary Results**). Our results are in line with the proposed model of TopR1/TopoVI-induced increase in positive DNA supercoiling upon heat stress (43, 44, 45) and adds that TopR1 is transcriptionally regulated in response to heat shock (**Figure 7b, Supplementary Results**). Given the absence of a dedicated heat-shock responsive transcription regulatory mechanism and the connection between chromatin organization and transcriptional activity in *Sulfolobales* (47), the observed dynamic regulation of DNA supercoiling and packaging could be hypothesized to play an important role in heat-shock responsive global regulation in Sulfolobales, at least on the transcriptional level.

## Discussion

In this work, we reveal that a temperature increase above the optimal growth temperature causes large changes in differential abundance in transcripts and proteins in *S. acidocaldarius*, with a fast and immediate response on the RNA level and a less pronounced and slower response on the protein level. Clearly, heat shock induces major effects on information processing, although it can be anticipated that this is not only the result of regulation taking place, but also of RNA and protein denaturation caused by a lack of thermostability and of changes in the synthesis and degradation rates (48), both of which are enzymatically catalyzed and thus inherently dependent on temperature. The use of a pulse-labeling approach enabled us to visualize RNA and protein neosynthesis and to demonstrate that global transcription and translation activity decreases in response to heat shock, albeit with different response dynamics.

The fast decline in transcriptional activity does not align with a lowered protein abundance of the basal transcription machinery consisting of the RNAP and general basal transcription factors. The exception to this observation is the initiation factor TFE, for which both subunits are significantly downregulated on the transcript and protein level. A heat-shock responsive depletion of TFE was previously also observed in the related species *S. solfataricus* (49). Notwithstanding temperature-dependent effects on the activity of all components of the basal transcription machinery, these results point to a key role for TFE, which optimizes transcription initiation by stabilizing the open transcription bubble (50, 51), by reducing the rate of transcription initiation upon heat stress. It could be hypothesized that the organism maintains a similar level of protein availability of other components of the transcription machinery to enable a fast recovery after the heat shock condition is relieved. In contrast to components of the transcription preinitiation complex, a considerable upregulation was observed on the protein level for TFS2 in response to heat shock. This transcription elongation factor is involved in RNA cleavage and restarting transcription within stalled elongation complexes (52).

In contrast to transcription, translational activity decreases slowly but consistently in response to heat shock. This is accompanied by a downregulation of translation initiation and elongation factors, charged tRNAs and ribosomes. Together with the finding that heat shock causes an immediate decrease in levels of neosynthesized 16S and 23S rRNA and the decrease in the abundance of many 30S and 50S ribosomal proteins, this demonstrates that there is a change in the translational capacity. In striking contrast, a considerable increase was observed for the expression of genes involved in tRNA and rRNA quality control. Whereas RNA-seq did not allow reliable quantification of tRNA, it has previously been shown that the tRNA pool was shifted upon stress conditions in yeast (53).

Despite lowered transcriptional and translational activities, the pulse-labeling experiments clearly indicated that neosynthesis is still taking place under the heat shock conditions chosen in these experiments. Consequently, it is likely that true regulatory events are taking place, especially for genes displaying a higher transcript and/or protein abundance upon heat shock. Changes at the transcriptomic level are not clearly correlated to those on the proteomic level; moreover, it is remarkable that despite the lower number of genes with a significantly lower or higher abundance at the protein level, the RNA level of many of these genes remained unchanged. Such a lack of correlation between transcriptomic and proteomic dynamics has also been observed in *S. acidocaldarius* for other stress conditions such as nutrient limitation (54) and was also found for heat shock response in bacteria (53, 54). It suggests a prevalence of post-transcriptional and/or post-translational regulatory events in addition to the extensive transcriptional regulation observed upon heat shock.

In contrast to eukaryotic and bacterial systems, as well as to the euryarchaeal *P. furiosus* and *A. fulgidus* (15, 16, 18), it seems unlikely that *S. acidocaldarius* harbors a classical heat shock transcription factor that is responsible for the transcriptional upregulation of important heat-shock responsive genes such as HSPs. This is corroborated by the absence of a conserved motif in heat shock regulon promoters and refutes previous hypotheses that heat-shock induced transcription factors might play such a role in *S. acidocaldarius* (20). Instead, a strong transcriptional upregulation might be mediated by TFB, and possibly TFE recruitment, given the highly conserved BRE and PPE signatures. Given the downregulation of TFE during heat shock, a strong PPE element could possibly discriminate between genes that require strong induction and the others.

Alternatively, it can be hypothesized that heat-shock induced effects on chromatinization and chromosome structuring underly the observed gene regulation on the transcriptional level. Indeed, our findings confirm the TopR1-model of increased DNA positive supercoiling upon heat shock (43, 44, 45). Furthermore, the chromosome of *S. acidocaldarius* and other Crenarchaeota is compacted by a large array of small bacterial-like NAPs (47). NAPs seem to play an important role in preventing DNA denaturation at heat-shock temperatures as observed and proposed for other hyperthermophilic archaea (57), however, they might also play a role in temperature-responsive global gene regulation. Previously, it was shown that cold shock leads to altered short-range and long-range interactions and an associated decreased ClsN occupancy and transcriptional activity in the chromosome of *S. islandicus* (58). Here, it is shown that heat shock induces altered levels of various chromatin proteins, not only ClsN, but also classical NAPs and Lrs14-family proteins, with the latter type also being involved in transcription regulation (41, 42, 46, 57). In conclusion, we hypothesize that in contrast to histone-harboring Euryarchaeota that have heat-shock transcription factors such as Phr, Sulfolobales and other histone-lacking thermophilic archaea employ an evolutionary ancient mechanism relying on temperature-responsive changes in DNA organization and compaction, induced by the action of NAPs and in combination with a considerable portion of post-transcriptional and -translational regulation.

## Supporting information

Supplementary Results; Supplementary Methods; Supplementary Figure; Supplementary Table

## Acknowledgements

We are grateful to Michael Jüttner and Nicolas Alexandre (SF-C lab, University of Regensburg) for their support in setting up *S. acidocaldarius* pulse-labeling experiments, Gaëtan Herinckx for technical assistance during mass spectrometry sample processing and Joris Van Lindt for help in troubleshooting the western blotting system on *S. acidocaldarius* crude lysate. We are grateful to Norio Kurosawa for the gift of the *S. acidocaldarius* SK-1 strain. This research was supported by Research Foundation Flanders (FWO-Vlaanderen) [PhD fellowship 1134419N to RB and Research Projects G021118N and G062820N to EP] and by the iBOF project “POSSIBL” [iBOF/21/092] of the Bijzonder Onderzoeksfonds. Research in the SF-C laboratory is generously supported by the German Research Foundation (DFG): individual research grant [FE1622/2-1] and collaborative research centre SFB/CRC 960 [grant SFB960-AP1, SFB960-B13] ‘RNP biogenesis: assembly of ribosomes and non-ribosomal RNPs and control of their function’. DG acknowledges funding by the German Research Foundation (grant number SFB/CRC 960 AP7).

