## Supplementary Results; Supplementary Methods; Supplementary Figure; Supplementary Table for "Transcriptional and translational dynamics underlying heat shock response in the thermophilic Crenarchaeon *Sulfolobus acidocaldarius*"

Rani Baes *et al*

#### Supplementary Methods

***Construction of the tagged thermosome strain***

With the aim of detecting and quantifying protein levels of individual thermosome subunits, a *Sulfolobus acidocaldarius* strain was constructed harboring C-terminally tagged thermosome subunits (Thα, *Saci_1401*; Thβ, *Saci_0666*; Thγ, *Saci_1203*) with each subunit gene fused to a different tag (FLAG-, His- or HA-tag, respectively). The SK-1*x*Thα-FLAG+Thβ-6xHis+Thγ-HA strain was constructed from SK-1 (1) based on the pop-in/pop-out strategy as previously described (2).

To this end, for each of the three subunit genes, genomic regions of about 1,000 base pairs (bp) up- and downstream of the intended C-terminal tag location were PCR-amplified using *S. acidocaldarius* MW001 genomic DNA as a template. Primers were designed to incorporate the desired tag. Subfragments representing the up- and downstream regions were fused in an overlap PCR approach and the resulting fragments with a length of about 2,000 bp were subsequently cloned in a BamHI/NdeI-digested pSVA431 plasmid vector, harboring *pyrEF* genes (2),using a SLiCE cloning strategy (3). *Escherichia coli* DH5α or MG1655 was used as a host for cloning and plasmid propagation, generating three different "pop-in" plasmids: pSVA431*x*Thα-FLAG, pSVA431*x*Thβ-6xHis and pSVA431xThγ-HA. All primers used in this work are presented in **Supplementary Table S2**, all plasmids in **Supplementary Table S3**.

*S. acidocaldarius* SK-1 electrocompetent cells were transformed by electroporation as described in (2) with the exception that the template plasmid was not methylated. For growth on plates, 0.6 % Gelrite® (Duchefa Biochemie, Netherlands, Haarlem) was used as a solidifying agent of Brock basal salts medium supplemented with 0.2 % sucrose, 0.1 % NZ-Amine, but lacking uracil, and acidified to pH 3.0-3.5 with $H_2SO_4$, with addition of 3 mM $CaCl_2$ and 10 mM $MgCl_2$. After incubating during 5 days at 75°C, colonies were treated by spraying with a solution of 5 mg $ml^{-1}$ 5-bromo-4-chloro-3-indolyl-b-D-galactopyranoside (X-gal), revealing "pop-in" integrants by blue color formation caused by LacS-activity. These colonies were further grown in liquid culture without uracil upon confirmation by PCR analysis. At mid/end-exponential phase ($OD_{600}$ of 0.6-0.8), cells were spread on plates containing 20 µg $ml^{-1}$ uracil and 200 µg $ml^{-1}$ 5-fluoroorotic acid (5-FOA) and incubated for 5 days at 75°C. Candidate "pop-out" transformants were inoculated in liquid medium with uracil, and presence of the desired C-terminal tag was confirmed by PCR analysis and Sanger sequencing. This procedure was performed first for the introduction of the Thβ-6xHis tag and repeated twice for introduction of the Thα-FLAG and Thγ-HA tag, respectively, finally generating a strain harboring all three tags in the genome (SK-1*x*Thα-FLAG+Thβ-6xHis+Thγ-HA).

***Analysis of cellular viability by plating***

Culture samples of SK-1*x*Thα-FLAG+Thβ-6xHis+Thγ-HA were diluted with Basic Brock medium to $OD_{600nm}$ 0.1 (*i.e.* the $10^{-1}$ dilution), based on the $OD_{600nm}$ measurement at the start of the experiment. A serial dilution series was constructed to $10^{-6}$ and 10 µL of each dilution were spotted on freshly prepared plates. For growth on plates, 0.6 % gelrite was used as a solidifying agent of the Brock medium with addition of 3 mM $CaCl_2$ and 10 mM $MgCl_2$. Plates were incubated for 5 days at 75°C and analyzed as previously described (4).

***Western blotting***

SK-1*x*Thα-FLAG+Thβ-6xHis+Thγ-HA cell pellets were resuspended in 500 µL lysis buffer (phosphate-buffered saline (PBS) pH 7.5, 1 % sodium dodecyl sulfate (SDS), 5 mM phenylmethylsulfonyl fluoride (PMSF), cOmplete™ Protease Inhibitor Cocktail (Roche, Switzerland, Basel)), lysed by ultrasonication (4 min at 70 %

amplitude with 1 min pulses at 4 °C) and centrifuged for 15 minutes at 16,100 *x g*. Total protein concentrations were determined employing the Pierce™ Rapid Gold BCA Protein Assay Kit (Pierce Biotechnology, Inc., USA, Rockford).Samples were normalized to equal concentration, mixed with 4x LDS loading dye, denatured for 10 minutes at 70°C and loaded four times (6.6 µg total protein per lane) on four separate but identical denaturing sodium dodecyl sulfate (SDS) polyacrylamide gel electrophoresis (PAGE) analyses (NuPAGE™ 4-12%, Bis-Tris, 1.0 mm Mini protein gel) and ran for about 35 minutes at 200 V. One gel was stained with Coomassie as loading control. Proteins from the three other gels were transferred to three Transblot Turbo 0.2 µm polyvinylidene fluoride (PVDF) membranes employing the Trans-Blot Turbo Transfer System with default settings for a mini gel. Membranes were blocked for 1 hour with PBS + 0.1% Tween-20 + 5% skimmed milk, after which the corresponding (primary) antibodies were added in fresh blocking buffer for overnight incubation at 4°C: Thβ-6xHis was targeted by monoclonal mouse anti-polyHistidine-HRP conjugate antibody (1:1000, A7058-1VL, Sigma-Aldrich, USA, Saint Louis), Thα-FLAG was targeted by monoclonal mouse anti-DYKDDDDK antibody (1:1000, 66008-3-Ig, ProteinTech, England, Manchester) and Thγ-HA was targeted by monoclonal mouse anti-HA antibody (1:500, 26183,Invitrogen, USA, Waltham). Blots for Thα-FLAG and Thγ-HA were washed twice for 30 minutes with PBS + 0.1% Tween-20 and subjected to secondary antibody binding for 1 hour at room temperature employing goat anti-mouse-HRP conjugate IgG (1:2000, SA00001-1, ProteinTech, USA, Waltham). All three blots were subsequently washed twice for 30 minutes with PBS + 0.1% Tween-20 and 30 minutes with PBS. Blots were developed by addition of HRP substrate (Pierce ECL Western Blotting Substrate, Thermo Fisher Scientific, USA, Eugene) and visualized with the Bio-Rad Gel Doc XR+ System (Bio-Rad, USA, Hercules). Blots were quantified by ImageJ 1.53e (5) and band signal intensities were standardized to the Coomassie signal of the complete lane.

***RNA sequencing***

Total RNA was extracted from MW001-stabilized cell pellets using the RNeasy Mini Kit (Qiagen, USA, Maryland) and on-column DNase treatment (Qiagen, USA, Maryland). Cell pellets were resuspended in 600 µL RLT™ lysis buffer and centrifuged for 10 minutes at 12,108 x *g* and 4°C. RNA was finally eluted from the column in 30 µL nuclease-free water. The total RNA quantity was determined with a Qubit RNA High Sensitivity Assay (Thermo Fisher Scientific, USA, Eugene) and RNA integrity was evaluated on a Bioanalyzer instrument with a RNA 6000 Nano chip (Agilent Technologies, USA, Santa Clara) (**Supplementary Dataset S1**). Ribosomal RNA depletion was established using a PAN-Archaea riboPOOL kit (siTOOLS, Germany, Planegg), followed by purification with a Zymo RNA Clean and Concentrator-5 kit (Zymo Research, USA, Irvine). Sequencing libraries were subsequently prepared with the TruSeq Stranded Total RNA Library Kit (Illumina, USA, San Diego) in combination with the RNA Unique Dual Indices (IDT for Illumina, USA, San Diego). Library enrichment PCR proceeded for 9 cycles. Library quality was verified on a Bioanalyzer instrument with a DNA High Sensitivity chip (Agilent Technologies, USA, Santa Clara) and concentrations were measured with qPCR according to 'Sequencing library quantification guide' (Illumina, USA, San Diego). Sequencing was performed on a NextSeq500 instrument (Illumina, USA, San Diego) in high output with single reads of 75 nts and 2% Phix spike-in (**Supplementary Dataset S2**).

Data were processed by first performing a quality control by FastQC (6); quality trimming and filtering was performed by Trimmomatic (7) (**Supplementary Dataset S2**). Sequencing reads were mapped to the *S. acidocaldarius* DSM639 genome (NC_007181.1) using STAR (8) and read counts were produced by RSEM (9). A minimum of 2182 genes out of 2351 coding genes were covered (= 92.8 %). Normalization and differential expression analysis was performed with the R-package EdgeR (10). Genes were considered differentially expressed if the false discovery rate (FDR) was lower than 0.05 (**Supplementary Dataset S3**).

***TMT-labeled Liquid Chromatography-Tandem-Mass-Spectrometry***

For protein extraction, cell pellets were resuspended in 500 µL lysis buffer (PBS pH 7.5, 1% SDS, 5 mM phenylmethylsulfonyl fluoride (PMSF), cOmplete™ Protease Inhibitor Cocktail (Roche, Switzerland, Basel), lysed by ultrasonication and centrifuged for 15 minutes at 16,100 *x g*. Total protein concentrations were determined employing the Pierce™ Rapid Gold BCA Protein Assay Kit (Pierce Biotechnology, Inc., USA, Rockford) (**Supplementary Dataset S1**). Samples of 450 µL lysate were flash-frozen and stored at - 80°C. Next, 100 µL lysate samples were diluted 1:10 with 30 mM triethylammonium bicarbonate (TEAB) followed by incubation at 95°C during 5 minutes. Trypsin was added at a 1:50 ratio followed by a 5-hour incubation at 37°C and a 10-minute centrifugation at 10,000 x *g* for the collection of peptide-containing supernatants. Digested samples were stored overnight at -20°C and peptides pelleted by speedvacevaporation. Pellets were resolved in 80 µL 30 mM TEAB using water bath sonication for 2 minutes. Remaining SDS was removed employing the Pierce™ Detergent Removal Spin Columns with a 0.5-ml volume (Pierce Biotechnology, Inc., USA, Rockford) using 30 mM TEAB as equilibration buffer. Peptides were quantified with the Pierce™ Quantitative Colorimetric Peptide kit (Pierce Biotechnology, Inc., USA, Rockford). For each peptide sample, 10 µg was Tandem Mass Tag (TMT) labeled using the TMTpro 16plex Label Reagent Set (Pierce Biotechnology, Inc., USA, Rockford). Samples were multiplexed to 120 µg, dried by speedvac centrifugation and the pellets were resuspended in 100 µl of 0.5 % trifluoroacetic acid in 5 % acetonitrile. Unincorporated TMT-labels were removed on a Pierce™ C18 Spin Column (Pierce Biotechnology, Inc., USA, Rockford) and eluted in 20 µL 70 % acetonitrile. Samples were dried by speedvac centrifugation and resuspended in 0.1 % trifluoroacetic acid 3.5% acetonitrile to a final peptide concentration of 0.5 µg/µL.

One µg of peptides were directly loaded onto a reversed-phase pre-column Acclaim PepMap 100 (Thermo Fisher Scientific, USA, Eugene) and eluted in backflush mode. Peptide separation was achieved using a reversed-phase analytical column Acclaim PepMap RSLC on an Ultimate 3000 RSLN nanoHPLC system (Thermo Fisher Scientific, USA, Eugene) as described by (11). Peptides were analyzed by mass spectrometry (MS) at an Orbitrap Fusion Lumos tribrid (Thermo Fisher Scientific, USA, Eugene) with enabled advanced peak determination (APD) and with relative quantification by MS2. Intact peptides were detected in the Orbitrap at a resolution of 120,000 with a scan range m/z from 375 to 1500 and an AGC target of $4x10^5$, maximum injection time was set to 50 ms. A data-dependent procedure of MS/MS scans was applied for the top precursor ions above a threshold ion count of $3.0 \times 10^4$ in the MS survey scan with 60 s dynamic exclusion. The total cycle time was set to 3 s. For MS2 quantification of the TMT reporter ions, MS/MS spectra were acquired in the Orbitrap at a resolution of 50,000 after HCD fragmentation at 35%, with an AGC target of $1 \times 10^5$ ions and a maximum injection time of 120 ms.

The resulting MS/MS data were processed using Sequest HT search engine within Proteome Discoverer 2.5 against the *S. acidocaldarius* reference target-decoy database obtained from NCBI (1 Jan 2022, 2,267 forward entries). Trypsin was specified as the cleavage enzyme, allowing up to two missed cleavages, four modifications per peptide, and up to three charges. Mass error was set to 10 ppm for precursor ions and 0.1 Da for fragment ions, and considered dynamic modifications were +15.99 Da for oxidized methionine and +42.011 Da for acetylation of the protein N-terminus. Fixed modifications were TMTpro (+304.207 Da) for lysine and peptide N-termini and +57.00 Da for carbamidomethyl cysteine. An overall of 1,115 proteins were detected out of the 2,267 predicted protein encoding genes (= 49.2 %). Differential protein expression analysis was performed using the DEqMS pipeline for TMT-labeled MS data (12). To this end, protein abundance values were $\log_2$ transformed, replicate outliers removed and data normalized to have equal medians in all samples. Benjamini-Hochberg corrected p-values (13) were considered statistically significant at a threshold < 0.05 (**Supplementary Dataset S3**).

#### Supplementary Results

##### *Heat shock proteins and proteasomal degradation*

Arguably the most detrimental effect of high temperature on the cell is the denaturation and aggregation of proteins. The classical set of heat shock proteins (HSPs) in *S. acidocaldarius* comprises a number of chaperones, which are involved in protein (re)folding at the optimal growth temperature and upon heat stress: the thermosome (HSP60), small HSPs (HSP14 and HSP20) and prefoldin (14). The thermosome complex, the major HSP in *Sulfolobus* (10, 11), is involved in ATP-dependent (re)folding of proteins (14) and has a different subunit composition (Thα, Thβ and Thγ) based on the temperature in an *in vitro* study (17). Even at 75°C, MS-MS data reveal that the thermosome subunits are the most abundant proteins in the cell (Thβ at position 1, Thγ at position 2 and Thα at position 4) (**Supplementary Figure S4**). We observed a rapid and considerable increase in gene expression for two thermosome subunits (15-60 minutes), *thα* and *thβ,* with a peak at 30 minutes after heat shock (**Figure 3**). This was reflected by a significant increase of the Thα and Thβ proteins 30 minutes after heat shock. In contrast, Thγ was not heat-shock responsive.

sHSPs (HSP14 and HSP20) and prefoldin have no intrinsic refolding abilities. Instead, they act as "holdases" by binding to aggregating proteins before transferring them to the thermosome complex for active refolding (14). Our results show a steady increase in HSP20 gene expression (15-60 minutes) and protein expression (30-60 minutes) (**Figure 3**), with HSP20 being most upregulated protein at 60 minutes after heat (**Supplementary Dataset S3**). Whereas no transcriptional regulation is taking place for *hsp14*, its protein level is increased after HS (60 minutes) (**Figure 3**), suggesting that HSP14 might be regulated on a post-transcriptional level. Prefoldin subunit α is transcriptionally downregulated (30 minutes) and subunit β is upregulated at 15 minutes at the RNA level and at 60 minutes at the protein level (**Figure 3**).

When the mis- or unfolded proteins cannot be rescued by the molecular chaperones, the cellular machinery for protein degradation takes over to prevent proteins aggregation. The 20S proteasome is an ATP-independent proteinase, generally degrading completely unfolded proteins (18), and is assisted by the proteasome-activating nucleotidase (PAN) $AAA^+$ ATPase and the proteasome assembly chaperone (PAC2). Post-translational modification of protein substrates by Urm1/SAMP, known as the archaeal-type urmylation or sampylation, can act as a signal for proteosome-mediated degradation, by direct recognition of the modified protein by the PAN-ATPase and the core proteasome *in vitro* (19). Indeed, a subset of transcripts involved in proteolytic degradation were differentially expressed upon heat shock (**Figure 3**). Whereas the mRNA level of the proteasome α subunit was not heat-shock responsive, a slight increase was observed for the β-subunits over the course of 60 minutes *post* heat shock. Contrary, we observed a steady decrease in transcripts encoding PAN ATPase and PAC2 (15-60 minutes). Moreover, we observed a considerable increase in Urm1 transcripts (15-60 minutes), associated with a slight decrease in transcripts encoding an ELSA homolog, a Urm1/SAMP activator (15-60 minutes). It is therefore possible that urmylation modifications are increased upon heat shock, although protein levels were not detected in our MS study.

Taken together that i) a substantial increased transcriptional gene expression was observed for the chaperone α and β thermosome subunits at 30 minutes after heat shock, followed by a slower increase in their protein levels at 30-60 minutes; ii) a lack of transcriptional upregulation was observed for proteolytic PAN ATPase and PAC2 at the 60 minute time point; iii) a 88°C heat shock did not impair cellular viability after 60 minutes, we can assume that the effect of the temperature stress for 60 min on the protein level is still manageable for the chaperones and does not (yet) require additional proteolytic support.

##### *DNA topology*

The array of DNA topoisomerases present in *S. acidocaldarius* consists of three type I topoisomerases (TopA and two reverse gyrases TopR1, TopR2), one type II topoisomerase (TopoVI) and all are involved in resolving topological stresses created by DNA-based processes (20). Whereas TopA levels remained undetectable in previous heat shock response studies in Sulfolobales (24, 25), we here show by RNA-seq that there is a strong decrease in *topA* transcripts immediately after heat shock, which is even further decreasing over the course of 60 minutes (**Figure 7b**). Even more, transcript counts of *topA* at 75°C are relatively high (average normalized cpm value of 291) and is about double the expression of *topR1* (cpm of 153) and one third of *topR2* (cpm of 608) (**Supplementary Dataset S3**). Given our observed downregulation of *topA* transcripts upon heat shock, the fact that TopA is efficient in DNA decatenation and relaxing negative supercoiling (23) (a topological state which is not prevalent upon heat shock), in combination with the observation that TopA is much less active at higher temperatures (22), our results provide evidence that TopA is not required in homeostatic control of DNA topology after heat shock.

The reverse gyrase enzyme has been described as a marker for thermophily and is able to introduce positive supercoiling in the DNA, which is thereby increasing DNA heat stability and preventing DNA damage (24). In our RNA-seq, transcript levels of *topR1* were decreased within 30 minutes of heat shock and *topR2* was shown not to be heat-shock responsive over the course of 60 minutes (**Figure 7b**). This timing is in perfect correspondence with the northern blotting observations of the *topR1* and *topR2*-transcripts in a *S. solfataricus* heat-shock experiment (21) and TopR2 western blotting (22). The two reverse gyrases of *S. acidocaldarius* are proposed to have different working mechanisms and sub-functionalizations *in vivo* (28, 29). TopR2 has been proposed to be the most important reverse gyrase at the optimal growth temperature, based on viability of deletion strains in *S. islandicus* (27), and has been proposed to be involved in DNA replication and repair (20). In contrast, TopR1 is involved in controlling DNA topology at high temperatures (20). Indeed, our results support the previous hypothesis of a TopR1-regulated model of supercoiling upon heat shock (22): the increased enzymatic activity of TopR1 at higher temperatures (25, 28) causes an increase in linking number (positive supercoiling) immediately after heat shock in *S. islandicus* (28). As shown in (25, 28), the number of TopR1 proteins is decreasing upon HS, which is associated with a gradual decrease in linking number and a new topological state of the DNA, adapted to the higher temperature after 60 minutes of heat shock (28). Based on our findings, we can conclude that TopR1 is transcriptionally regulated in response to HS in *Sulfolobus*. However, this effect is not noticeable at the protein level in our study.

Lastly, the two subunits of TopoVI, a type II topoisomerase able to relax both negative and positive supercoils at the optimal growth temperature by cleaving double-stranded DNA (20). TopoVI subunit A was transcriptionally upregulated immediately after heat shock and transcript levels are continuing to increase over the course of 60 minutes (**Figure 7b**). TopoVI subunit B was also upregulated at the protein level at 60 minutes. A previous study has shown that TopoVI B is the limiting subunit in functional TopoVI and that TopoVI is only active with positively supercoiled substrate at 88°C, but activity is low (22). It is therefore reasonable to assume that TopoVI needs to increase in abundance to fulfill its role in decreasing the linking number of the positively supercoiled DNA after about 60 min (28) as a result of increased TopR1 activity (22). Indeed, our results suggest that more functional TopoVI is present after 60 min of heat shock.

##### *Motility and biofilm formation*

Heat shock has considerable effects on the expression of the major cellular appendages, the adhesive type IV pili (aap pili) and the archaellum, and their corresponding regulators (**Supplementary Figure S5**). Whereas the aap pili are indispensable for surface adhesion and biofilm formation (29), rotary motion of the archaellum drives cell motility (30). The pilus consists of AapA and AapB pilins (29), which showed a different trend of expression upon heat shock (15-60 minutes): whereas *aapA* expression was strongly upregulated, *aapB* was strongly downregulated (**Supplementary Figure S5**). The motor complex anchor protein *aapE* was downregulated (15-60 minutes) and *aapF* was strongly upregulated (15-60 minutes). However, both motor proteins had decreased translational levels. Upon heat shock, transcript levels of the archaellin filament (*arlB*) remained unchanged, however, we observed a transcriptional downregulation of the a*rlX* scaffold (30 minutes) and upregulation of the *arlG* and *arlJ* motor (**Supplementary Figure S5**).

Regulation of motility and biofilm formation, as well as the switch between a free-living, motile, and sessile life style, is highly regulated, and depends on many regulators and their phosphorylation status (20, 21, 22). Our results demonstrated that all known regulators are differentially expressed upon HS (**Supplementary Figure S5**). Besides their role as nucleoid-associated proteins (NAPs), Lrs14-type proteins AbfR1 and Saci_1223 are key players in regulation of these physiological processes: AbfR1 activates motility, while repressing biofilm formation, and Saci_1223 activates biofilm formation (31). Upon heat shock, AbfR1 was strongly upregulated at the RNA (15-60) and protein levels (60 minutes). In addition, RNA levels of the archaellum activators *arnR* and *arnR1* activators were increased (60 minutes), while the RNA levels of the ArnS archaellum activator were found to be decreased (15, 60 minutes). In addition, we observed a fast, persistent transcriptional downregulation of the archaellum repressors ArnA and ArnB, kinase ArnD and phosphatase PP2A (15-60 minutes) upon heat shock. The kinase *arnC* was only transiently downregulated (15 minutes).

At optimal growth conditions in rich, liquid medium, there is typically no archaellum expression (32). However, previous studies have shown that motility is enhanced when *S. acidocaldarius* cells are exposed to high temperatures (34) or upon nutrient starvation (32). The ArnS activator plays an important role in starvation-induced motility (32). Here we show that in contrast, heat-shock induced motility is most likely not dependent on ArnS, given the lack of transcriptional upregulation observed (**Supplementary Figure S5**).

***DNA replication, genome segregation and cell division***

In response to heat shock, considerable differential expression related to DNA replication and cell division was observed. Replication in *Sulfolobus* is initiated during the S-phase of the cell cycle at three independent replication origins (*oriC1-3*). The major regulatory step in replication initiation consists of binding of a replication initiator protein to its corresponding origin(s) (35). Our RNA-seq and MS experiments showed that the initiator protein Cdc6-1 and Crenarchaeal-specific Whip are not heat-shock responsive over the course of 60 minutes (**Supplementary Figure S6a**). In contrast, Cdc6-3 was transcriptionally downregulated immediately after heat shock (15-60 minutes) and Cdc6-2 translationally downregulated (60 minutes). A previous study hypothesized that Cdc6-1 and Cdc6-3 are involved in promoting replication, given their main presence in the $G_1$ and S phase, whereas Cdc6-2 may be a negative regulator for replication, since it is only present in $G_2$ (36). It is therefore reasonable that replication initiation is not enhanced and it is even plausible that replication initiation is downregulated upon heat shock in *S. acidocaldarius*. Moreover, given the specificity of the initiator proteins (35) and the observation that Cdc6-2 is an essential transcriptional activator for upregulation of double strand break (DSB) repair proteins (see below) (37), it is possible that firing of the different origins and activation of DSB repair is affected upon heat shock.

Cdc6 binding is followed by the recruitment of the replication elongation machinery. Upon heat shock, a transcriptional downregulation was apparent for many proteins involved (**Supplementary Figure S6a**). We observed a decreased gene expression for the Mcm DNA helicase (at 60 minutes) and a considerable transcriptional downregulation of the single-strand DNA binding protein (SSB) immediately after the temperature shift (15-60 minutes). Whereas the large subunit of replication factor C (Rfc) and proliferating cell nuclear antigen (PCNA) subunits A and B were not heat-shock responsive, the small Rfc and PCNA subunit C were strongly transcriptionally downregulated (15-60 minutes) and slightly increased at the protein level (60 minutes). Transcript levels of the main replicative DNA polymerase *polB1* were transiently decreased (15 minutes), associated with a decrease of PolB1-binding protein PBP-1, but not PBP-2. In addition, *fen1* and *lig1* were transcriptionally downregulated upon heat shock, and decreased PriL and Fen1 protein levels were noticeable at 60 minutes. A persistent transcriptional downregulation was observed for *gins23* (30-60 minutes). Taken together, this might suggest that the overall process of DNA replication is slowed down in response to heat shock.

Daughter chromosomes remain in close contact during the extensive $G_2$-phase, which is hypothesized to facilitate homologous recombination (HR)-mediated repair of DNA when exposed to harsh environmental growth conditions (38). Although the exact mechanism is still under study, subsequent genome segregation is mediated by SegAB (27, 28) (**Supplementary Figure S6b**). SegB transcripts levels were unaffected by heat shock, yet, protein levels were slightly increased (60 minutes). In contrast, SegA was strongly transcriptionally (15-60 minutes) and translationally (60 minutes) downregulated. Although SegA polymerization is enhanced by SegB, the decreased SegA levels suggest that less functional segregation machinery can form after heat shock.

Cell division in *S. acidocaldarius* is mediated by a set of Cdv proteins (**Supplementary Figure S6b**). Upon heat shock, all Cdv-encoding transcripts show a significant, continuous transcriptional downregulation (15-60 minutes). In combination with the observation that growth is impaired in the *ΔcdvB1*, *ΔcdvB2* and *ΔcdvB3* mutants (29, 30), this makes it unlikely that cell division will ensue upon heat shock. This is further supported by the slight transcriptional decrease in proteasomal assembly and accessory proteins that was observed upon heat shock (**Figure 3**). Moreover, the genes encoding the *Sulfobus*' S-layer, *slaA* and *slaB*, are both transcriptionally downregulated upon heat shock (**Supplementary Figure S6b**). However, SlaA protein is slightly increased in abundance at 60 minutes.

Our study suggests that all steps of DNA replication, DNA segregation, cell division and S-layer synthesis are slowed down or halted upon heat shock. It is possible that reduction of cell division might limit genome segregation. Thus, it is plausible to assume that heat shock arrests the cells in $G_2$ phase and induces a cell division block, similar to cells that enter stationary phase (42).

***DNA repair and DNA import***

Increasing the positively supercoiled topology of the DNA upon heat shock by the action of TopR1 is *in se* an important way to prevent DNA damage, given that this prevents formation of single-stranded (ss) DNA, which is more prone to degradation. In contrast, DNA-based processes also require opening of the double-stranded (ds) helix. Given the observed downregulation of the ssDNA-protecting protein SSB (**Supplementary Figure S6a**) and the fact that SBB enhances reverse gyrase activity in a TopR1/TopR2 mix in *Sulfolobus sp*. (43), it is possible that SBB and TopR1 are jointly acting in the homeostatic control of DNA supercoiling and decrease of DNA replication upon heat shock.

Besides the main replicative DNA polymerase PolB1, *S. acidocaldarius* encodes three other accessory DNA polymerases (PolB2, PolB3 and PolY), of which the exact functions are not yet thoroughly understood, besides that these are mainly involved in DNA repair and DNA damage tolerance (33, 34). Upon heat shock, a decrease of *polB2* transcript levels (60 minutes) was observed as well as an immediate transcriptional upregulation of *polB3* and *polY* (15-60 minutes) (**Supplementary Figure S7a**). A previous study showed that *S. acidocaldarius* accessory DNA polymerase single, double and triple deletion strains are more sensitive to heat shock than the wild-type strain, especially *ΔpolB3* strains (45). Furthermore, PCNA stimulates polymerase activity of PolY, which is able to bypass DNA damage, and reduces the rate of single base deletions (46). Given the differential expression of PCNA-C upon heat shock (**Supplementary Figure S7a**), PolY activity could be affected as well. Moreover, it was previously shown that *S. solfataricus* PolY is inhibited by TopR1 and directly or indirectly stimulated by SBB, thereby preventing introduction of mutations and increasing genome stability (44). Our results thus suggest a crucial role for PolB3 and PolY in controlling DNA stability upon heat shock.

Despite adaptation of the DNA topology, spontaneous DNA damage increases at elevated temperatures *e.g.*: hydrolytic deamination and depurination, oxidation of guanines, strand breakage,… (47). The need for repair of such DNA damage is crucial in order to allow for DNA replication and maintain genome stability (48). In archaea, four universal DNA repair pathways are conserved (47), and upon heat shock, considerable changes were observed in expression levels of many enzymes involved (**Supplementary Figure S7a**). In *Pyrococcus furiosus*, the mismatch-specific endonuclease EndoMS (NucS) has been identified to be the key player correction of mismatched bases incorporated during DNA replication by mismatch repair (MMR) (50, 52). In *S. acidocaldarius*, we found that *endoMS* is not heat-shock responsive (**Supplementary Figure S7a**), which is coherent with the observation that DNA replication is decreased upon heat shock (see above). Structural perturbations of the DNA helix as a result of photoproduct-induced lesions are removed by nucleotide excision repair (NER) (47). The presence and exact working mechanism of NER in *Sulfolobus* is still speculative, nevertheless, putative enzymes involved are differentially expressed at the transcriptional level upon heat shock, including XPD helicase, XPB paralog *XPB1*, but not *XPB2*, and *Bax1* (**Supplementary Figure S7a**). The most prevalent type of DNA damage is caused by hydrolytic depurination, deamination of cytosine, oxidation or methylation and is corrected by base excision repair (BER). Depurination is easily triggered by reactive oxygen species and the rate of deamination is increasing at higher temperatures, possibly changing base-paring propensities and increasing mutation-rate (50, 53). Damaged bases are detected and the glycosidic bond cleaved either by a glycosylase specific for the damage base and AP endonuclease Endo III (54, 55) or endonucleases such as endoV (47). Additional processing might occur by flap displacement, excision by the Fen1 nuclease (50, 53) and PCNA (53). Upon HS, we observe a transcriptional downregulation of type IV DNA glycosylase, *endo III*, *fen1* and DNA ligase and an upregulation of *endoV* (**Supplementary Figure S7a**).

The main type of DNA damage is occurring from DNA double strand breaks (DSBs), which block DNA-based processes and can induce major mutations, genome rearrangements and cell death (47). Given the absence of the non-homologous end joining pathway in archaea, the repair pathway is homologous recombination, which is dependent on the presence of a second intact copy of the DNA (50, 57). Besides the use of homologous recombination in DSB repair, it is also involved in restarting DNA replication at stalled forks (47). Upon HS, we noted a considerable downregulation of enzymes involved in DSB end recession: gene expression of *mre11*, *rad50* (15-60 minutes) and *nurA* (15-30 minutes), and protein levels of Mre11 (30-60 minutes) and NurA (60 minutes) (**Supplementary Figure S7a**). Whereas HerA transcript levels were not heat-shock responsive, protein levels were considerably increased (60 minutes). RadA recombinase was upregulated at the RNA (15-60 minutes) and protein level (60 minutes). Two enzymes involved in strand exchange were transcriptionally downregulated: helicase *hel308* (15 minutes) and Holliday junction resolvase *hjr saci_1741* (15-60 minutes). However, *hjr Saci_1558* was upregulated (60 minutes). The response upon heat shock is different than the considerable transcriptional upregulation of DSB repair genes observed for *S. solfataricus* upon exposure to ionizing radiation (55), and a moderate induction upon UV stress (56). Taken together, our results suggest a decreased need for DSB end recession upon high temperature stress. However, given the upregulation of RadA (**Supplementary Figure S7a**) and downregulation of SBB in the identical time-frames (**Supplementary Figure S6a**) and a previous finding that *S. solfataricus* SSB inhibits the RadA ssDNA dependent ATPase activity (57), it is plausible to assume that RadA has a key role in DSB repair upon heat shock.

Most of the time a second copy of the *S. acidocaldarius* chromosome is present in the cell, since most cells are in $G_2$-phase (38) and DNA replication and cell division is slowed down upon HS. However, it has been previously described for *S. acidocaldarius* that, upon exposure to UV irradiation, the UV-inducible (ups) pili and the Crenarchaeal system for exchange of DNA (Ced) are transcriptionally upregulated (56) and are involved in species-specific aggregation and chromosomal DNA import, respectively. This DNA copy is serving as a template for the repair of DSBs by homologous recombination (57, 61, 62, 63). In our study, and in contrast to UV stress (56), we observe no massive transcriptional upregulation of the ups-genes upon heat shock (**Supplementary Figure S7b**). However, a large transcriptional upregulation is observed for *cedA*, *cedA1*, *cedA2* and the HerA-homolog immediately after heat shock (15-60 min), associated with a downregulation of *cedB* shock (**Supplementary Figure S7b**). A transcriptional upregulation is observed for most of the DNA-processing enzymes encoded downstream of *ups*, which are thought to be involved in subsequent DNA processing (54), including *endo III* nuclease, a glycosyltransferase and the helicase Hel112, and a translational upregulation of the ParB-like nuclease.

Upon UV stress, it has been shown previously that the transcription initiation factor TFB3 is upregulated at 45 min and is serving as an essential activator for *ups*- and *ced* transcription at 90 min (61). Unfortunately, read counts for TFB3 were low in our RNA-Seq or MS analysis even at 75°C, so it remains to be analyzed whether TFB3 is required in the observed HS-responsive upregulation of *ced* gene expression. However, given that the upregulation of *ced* is already observed at 15 min of HS, it seems unlikely that this is dependent on prior upregulation of TFB3 (given that abundance is low at 75°C). Thus, our results suggests that, besides the second ‘own’ genomic copy, there is an additional need for intact DNA copies to enhance the repair the DSB induced by HS in *Sulfolobus*, but possibly haploid Crenarchaea in general. Alternatively, this might pose a way to increase genetic diversity upon temperature stress.

***Post-transcriptional and post-translational modifications***

In order to rapidly modify activity, function and stability of an RNA species or protein, a universally conserved mechanism consists of the addition of post-transcriptional and or post-translational modifications (62). It has previously been shown that ribose methylation of rRNA and $N^4$-acetylcytidine modification of RNA increases at higher cultivation temperatures of archaea, which might play a role in structural stabilization (66, 67).

Of all post-translational modifications, reversible protein phosphorylation is the most prevalent in *S. acidocaldarius* and is occurring on serine, threonine and tyrosine amino acid residues (65). Upon heat shock, we observed that many kinases are differentially expressed upon heat shock (**Supplementary Dataset S3**). Furthermore, there was an immediate transcriptional upregulation of the phosphatases *Saci-PTP* (15 minutes) and a downregulation of *Saci-PP2A* (15-60 minutes). A variety of protein kinases and phosphorylated transcriptional regulators have been characterized thus far in *Sulfolobales spp.* involved in a wide range of cellular processes (69, 70), including: chromosome organization, motility by the archaellum and biofilm formation (35, 71), fatty acid metabolism (69), DSR by homologous recombination (70),… The two protein phosphatases in *S. acidocaldarius* also have each their unique dephosphorylation target residues and profile (65). Therefore, small changes in phosphorylation status upon heat shock, could have major consequences on all different cellular processes.

Methylation on lysine residues is also a prevalent posttranslational modification in thermophiles as a strategy to increase protein thermostability and infer another layer of regulation (71). Methylations occur on proteins belonging to all arCOG categories, including many chromatin-associated proteins in *S. islandicus* (such as Alba-1, Alba-2, Cren7 and the Sac7d homolog) (71) and DNA replication and transcription associated proteins. Methionine adenosyltransferase, involved in the generation of the major methyl donor, is transcriptionally downregulated upon heat shock (15-60 minutes), yet, shows increased protein levels (60 minutes) (**Supplementary Dataset S3**). A lysine methyltransferase (*Saci_1539*) is transcriptionally downregulated (15-60 minutes).

Furthermore, glycosylation of proteins, *e.g.* the S-layer subunit SlaA, contributes to thermal stability by limiting the peptide backbone flexibility (62). Many enzymes involved in this pathway were increased in abundance upon heat shock (**Supplementary Dataset S3**) *e.g.* oligosaccharyl transferase aglB (RNA, 15-60 minutes), archaeal glycosylation enzyme 3 (protein, 60 minutes) and a predicted phosphoglucomutase/phosphor mannomutase and predicted 3-hexulose-6-phosphate synthase (protein, 60 minutes). Thus, besides the observed response at the transcriptome and proteome level, heat shock induces major changes at the post-transcriptional and post-translational level.

### Supplementary Tables

##### Supplementary Table S1. Microbial strains used in this study.

| Name | Description | Reference |
|---|---|---|
| *S. acidocaldarius* DSM639 | *S. acidocaldarius* wild-type strain | DSMZ |
| *S. acidocaldarius* MW001 | Uracil-auxotrophic (*ΔpyrEF*) strain | (2) |
| *S. acidocaldarius* SK-1 | Uracil-auxotrophic (*ΔpyrEF*) and SuaI restriction system-deficient (*ΔsuaI*) strain | (1) |
| *S. acidocaldarius* SK-1*x*Thβ-6xHis | SK-1 expressing thermosome β (Saci0666) fused to a C-terminal 6xHis-tag (HHHHHH) | This study |
| *S. acidocaldarius* SK-1*x*Thα-FLAG+Thβ-6xHis | SK-1 expressing thermosome α (Saci1401) fused to a C-terminal FLAG-tag (DYKDDDDK) and thermosome β (Saci_0666) fused to a C-terminal 6xHis-tag (HHHHHH) | This study |
| *S. acidocaldarius* SK-1*x*Thα-FLAG+Thβ-6xHis+Thγ-HA | SK-1 expressing thermosome α (Saci1401) fused to a C-terminal FLAG-tag (DYKDDDDK), thermosome β (Saci0666) fused to a C-terminal 6xHis-tag (HHHHHH) and thermosome γ (Saci1203) fused to a C-terminal HA-tag (YPYDVPDYA) | This study |
| *E. coli* DH5α | Strain for plasmid cloning and propagation | Gibco |
| *E. coli* MG1655 | Strain for plasmid cloning and propagation | Gibco |

**Supplementary Table S2. DNA oligonucleotides used in this work for the construction of SK-1 (derivative) strains expressing tagged thermosome subunits.** FW = forward; RV = reverse.

| Name | Description | Sequence (5' → 3') |
|---|---|---|
| RB134 | FW subfragment PCR 1,000 bp upstream Thα-FLAG = FW overlap PCR | CTCAAGCTATGCATCCAACGCGTTGGGAGCTCTCCCATATGGTGCTAGACAAAGAAGTAGTACATGCAG |
| RB135 | RV subfragment PCR 1,000 bp upstream Thα-**FLAG** | GAGAGAGAATGTAAAATAATAAAAAAATAATATACTGTATTCA**TTTATCATCATCATCTTTATAATC**CTCTAATGAAGGTGCGCCAGGAG |
| RB136 | FW subfragment PCR 1,000 bp downstream Thα-**FLAG** | CTCCTGGCGCACCTTCATTAGAG**GATTATAAAGATGATGATGATAAA**TGAATACAGTATATTATTTTTTATTATTTTACATTCTCTCTCATTTATACATC |
| RB137 | RV subfragment PCR 1,000 bp downstream Thα-FLAG = RV overlap PCR | GAGCCAAGTACTAGAACTGCTCAAACCTAGGTCAGGATCCGAAAAGGGATTAGTTGCGCATTG |
| RB037 | FW subfragment PCR 1,000 bp upstream Thβ-6xHis = FW overlap PCR | CTCAAGCTATGCATCCAACGCGTTGGGAGCTCTCCCATAACAAATAGTGTATGGAATTATAGTTGATAAAGAAGTAG |
| RB038 | RV subfragment PCR 1,000 bp upstream Thβ-**6xHis** | GATCGCACAGTATAATAAAGGTAAAAAAAGTTACTTA**GTGGTGGTGGTGGTGGTG**GTCTTCTTCTTTACCTTTTTCAGACTCTTTC |
| RB039 | FW subfragment PCR 1,000 bp downstream Thβ-**6xHis** | GAAAGAGTCTGAAAAAGGTAAAGAAGAAGAC**CACCACCACCACCACCAC**TAAGTAACTTTTTTTACCTTTATTATACTGTGCGATC |
| RB040 | RV subfragment PCR 1,000 bp downstream Thβ-6xHis = RV overlap PCR | GAGCCAAGTACTAGAACTGCTCAAACCTAGGTCAGGATCATCATGACATAGAATTAGGGGAAGTTATAAATAATTATG |
| RB138 | FW subfragment PCR 1,000 bp upstream Thγ-HA = FW overlap PCR | CTCAAGCTATGCATCCAACGCGTTGGGAGCTCTCCCATATGGAGAAAGCGTAGACGAGACAACTTTAG |
| RB139 | RV subfragment PCR 1,000 bp upstream Thγ-**HA** | CATGAGGGAAAGAAAAGAAATCCTATATTTTAACTTTTTA**TGCATAATCAGGTACATCATAAGGATA**TCCCATAGGATATTGAGGCATTTGTTG |
| RB140 | FW subfragment PCR 1,000 bp downstream Thγ-**HA** | CAACAAATGCCTCAATATCCTATGGGA**TATCCTTATGATGTACCTGATTATGCA**TAAAAAGTTAAAATATAGGATTTCTTTTCTTTCCCTCATG |
| RB141 | RV subfragment PCR 1,000 bp downstream Thγ-HA = RV overlap PCR | GAGCCAAGTACTAGAACTGCTCAAACCTAGGTCAGGATCCGAACCCCGGTAGTTGCATACTC |
| EP393 | FW colony PCR cloning into pSVA431 | ATGACCATGATTACGCCAAG |
| EP394 | RV colony PCR cloning into pSVA431 | TCGAACTTGCAGACAAGTTC |
| RB144 | FW sequencing pSVA431xThα-FLAG (binding in 1,000 bp upstream) | GTGCAGTCGAGTCAGAGTTAG |

| | | |
|---|---|---|
| RB145 | RV sequencing pSVA431xThα-FLAG (binding in 1,000 bp downstream) | CTACTCATAGACGACATGAACCTAC |
| RB045 | FW sequencing pSVA431x Thβ-6xHis (binding in 1,000 bp upstream) | GTGGCACGGAATAAATGTATATAC |
| RB047 | RV sequencing pSVA431x Thβ-6xHis (binding in 1,000 bp downstream) | CATAATAACGCAGTCTCCTTCTAGTATAG |
| RB146 | FW sequencing pSVA431x Thγ-HA (binding in 1,000 bp upstream) | GAACGTTATAGAGAGCCCATACA |
| RB147 | RV sequencing pSVA431x Thγ-HA (binding in 1,000 bp downstream) | TGATCGACGTAAAGTATACCAAAGAC |
| RB158 | FW colony PCR/sequencing "pop-in" pSVA431xThα-FLAG | CACCTTCATTAGAGGATTATAAAGATGATGATGATAAATGAATACA |
| RB061 | FW colony PCR/sequencing "pop-in" pSVA431xThβ-6xHis | GAAGACCACCACCACCACCACCACTAAG |
| RB160 | FW colony PCR/sequencing "pop-in" pSVA431xThγ-HA | CTATGGGATATCCTTATGATGTACCTGATTATGC |
| RB062 | RV colony PCR/sequencing "pop-in" pSVA431(-derivatives) (binding pyrEF in pSVA431 backbone) | GATGACTACTTTAGAATATTCGAACTTGCAGACAAGTTCTATG |
| RB219 | FW colony PCR/sequencing "pop-out": Thα-FLAG | CTCAGGTACTAAAGAGTGCTGTAGAG |
| RB220 | RV colony PCR/sequencing "pop-out": Thα-FLAG | CTCAGTCCAGATTTTATCAACCTAGTTTTC |
| RB045 | FW colony PCR/sequencing "pop-out": Thβ-6xHis | GTGGCACGGAATAAATGTATATAC |
| RB047 | RV colony PCR/sequencing "pop-out": Thβ-6xHis | CATAATAACGCAGTCTCCTTCTAGTATAG |
| RB221 | FW colony PCR/sequencing "pop-out": Thγ-HA | GAAGACGTGACCAAGGAGAACATC |
| RB222 | RV colony PCR/sequencing "pop-out": Thγ-HA | GTGCCAAATGGTTAACGGCAATAC |

**Supplementary Table S3. Plasmids used in this work for construction of SK-1 (derivative) strains expressing tagged thermosome subunits.** Nucleotide sequences can be provided upon request.

| Name | Description | Reference |
|---|---|---|
| pSVA431 | Plasmid vector | (2) |
| pSVA431*x*Thα-FLAG | Pop-in/pop-out plasmid for fusing thermosome α (*Saci1401*) to a C-terminal FLAG tag (DYKDDDDK). | This work |
| pSVA431*x*Thβ-6xHis | Pop-in/pop-out plasmid for fusing thermosome β (*Saci0666*) to a C-terminal 6xHis tag (HHHHHH). | This work |
| pSVA431*x*Thγ-HA | Pop-in/pop-out plasmid for fusing thermosome γ (*Saci1203*) to a C-terminal HA tag (YPYDVPDYA). | This work |

#### Supplementary Figures

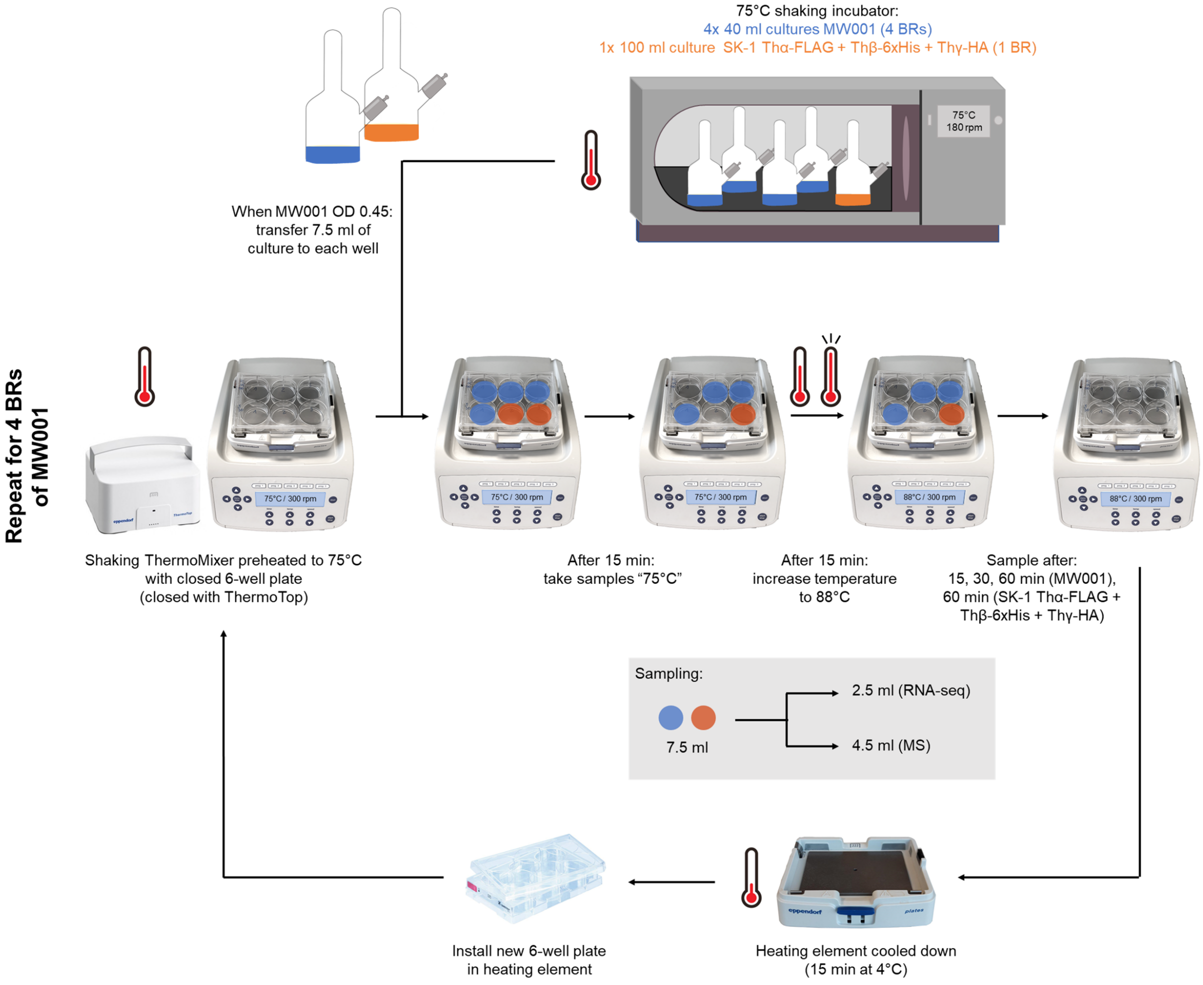


**Supplementary Figure S1. Heat-shock sampling for RNA-seq and MS.** Four separate *S. acidocaldarius* MW001 cultures, biological replicates (BRs), of 40 mL were incubated at 75°C until mid-exponential phase ($OD_{600nm}$ of ± 0.45). Then, 7.5 ml of each MW001 culture (BR) at a time was quickly transferred to each of four wells of a 6-well plate installed in a shaking 75°C heating block (ThermoMixer C® (Eppendorf, USA, Enfield), closed with ThermoTop® (Eppendorf, USA, Enfield)to avoid evaporation and condensation of the cultures). In each of the two remaining wells, 7.5 mL of a SK-1xThα-FLAG+Thβ-6xHis+Thγ-HA culture was added ($OD_{600nm}$ 0.17-0.35). Cultures were left to incubate for 15 minutes at 75°C, after which a sample of MW001 and SK-1*x*Thα-FLAG+Thβ-6xHis+Thγ-HA ("75°C samples") was collected. Then, 15 minutes after sampling, the temperature of the heating block was shifted from 75°C to 88°C (taking about 4.5 minutes) to trigger a heat shock response. The moment that the heating block itself reached 88°C corresponds to time point zero. It is important to note that i) the temperature inside the cultures is still increasing until a steady heat shock-temperature is reached after about 20 minutes, ii) the maximum, steady heat shock temperature that was reached inside the culture corresponds to 86°C and will be referred to during this study as such (**Figure 1a**). For MW001, samples were taken after 15, 30 and 60 minutes of heat shock administration, while SK-1*x*Thα-FLAG+Thβ-6xHis+Thγ-HA was sampled at 60 minutes only. The culture samples were split for transcriptomic (2.5 ml) and proteomic (4.5 ml) analysis. Although identical samples were taken for SK-1*x*Thα-FLAG+Thβ-6xHis+Thγ-HA, these were not processed for omics analyses. After sampling was complete, the heating element was removed from the heating block and cooled down (15 minutes at 4°C) to speed up the cooling process. A new 6-well plate was installed in the heating element, which was again preheated to 75°C in the heating block. This process was repeated consecutively for a total of four BRs of MW001, each time incorporating the same SK-1*x*Thα-FLAG+Thβ-6xHis+Thγ-HA as the control strain. All cell pellets were stored at – 80°C until further processing. More detailed information about the cultures and samples can be found in **Supplementary Dataset S1**.

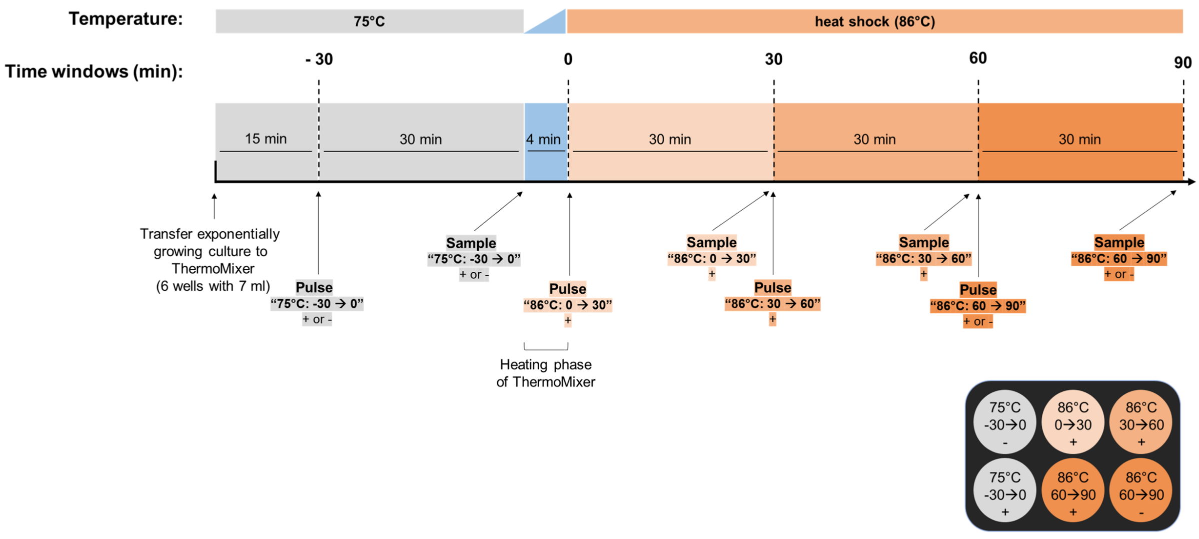


**Supplementary Figure S2. Experimental design of pulse-labeling experiments.** Pulse labeling was performed for neosynthesized RNA and protein at 75°C and upon a heat shock of 86°C. Cultures were grown until mid-exponential phase at 75°C, after which 7 ml of culture was transferred to 6 wells in a preheated thermomixer and left to incubate for 15 minutes. An excess of pulsing reagent (+) and a mock-control (-) were added to a well (scheme on bottom right) and left to incubate for 30 min at 75°C (samples 75°C: -30→0, +/-). After sampling the pre-heat shock sample, a 86°C heat shock was triggered as described in **Figure 1a**. Different heat shock time windows were pulsed with + (86°C: 0→30 / 30→ 60 / 60→90, +). One mock-control was incorporated during heat shock at the last time window (86°C: 60→90, -).

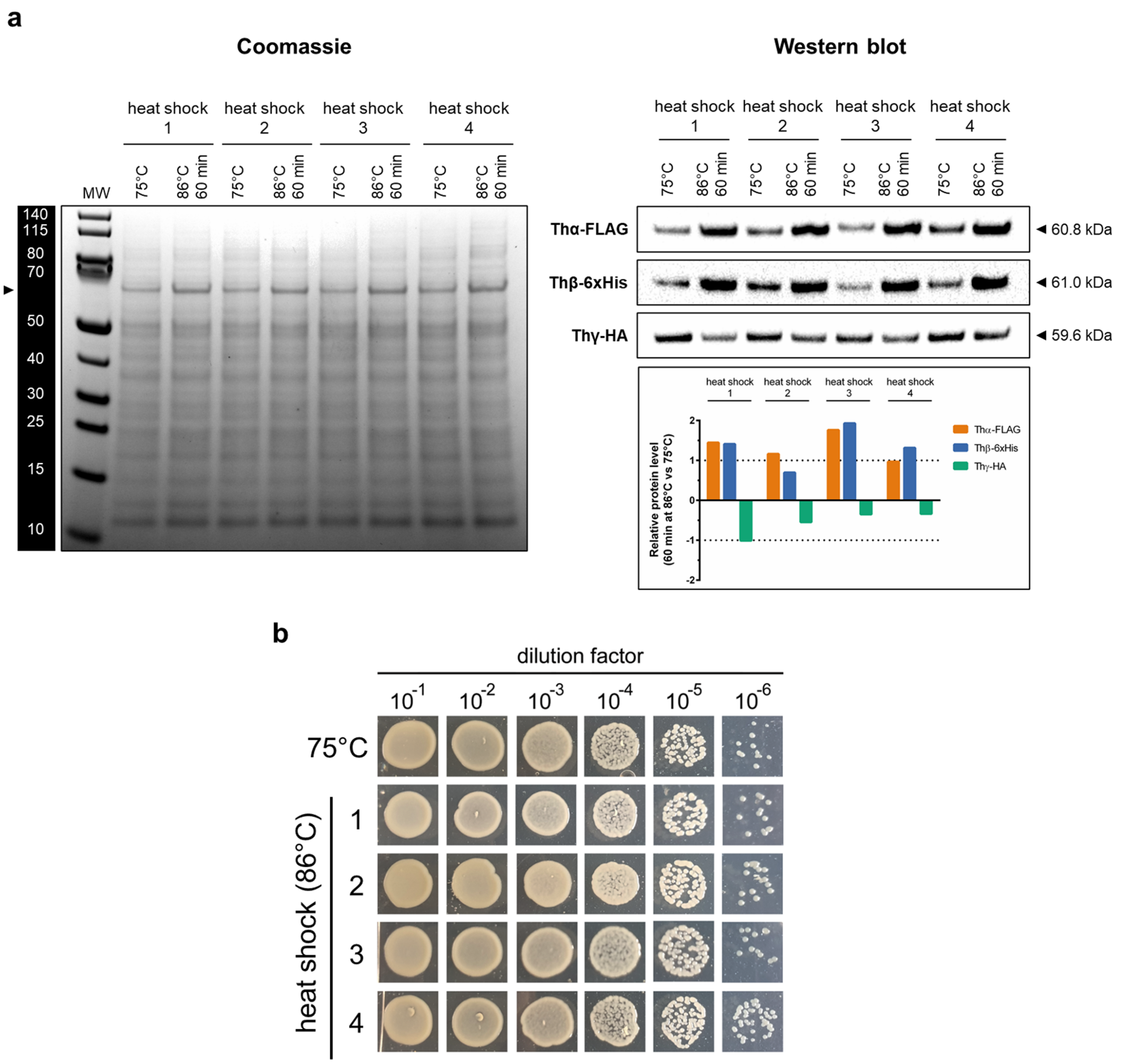


**Supplementary Figure S3. Validation of induction of heat shock response in the SK-1 Thα-FLAG + Thβ-6xHis + Thγ-HA control strain in the heat shock set-up of RNA-seq and MS. a** Western blot of the three thermosome subunits Thα-FLAG, Thβ-6xHis and Thγ-HA at 75°C and after 60 min at 86°C. Cell lysates (6.6 µg total protein per sample) were analyzed on four separate denaturing SDS-PAGEs. One gel was stained with Coomassie as loading control (left panel). Proteins from the three other gels were transferred to three Transblot Turbo 0.2 µm PVDF membranes employing the Trans-Blot Turbo Transfer System. One specific thermosome subunit was targeted per membrane: Thβ-6xHis was targeted by monoclonal mouse anti-polyHistidine-HRP conjugate antibody. Thα-FLAG and Thγ-HA were targeted by monoclonal mouse anti-DYKDDDDK antibody (1:1000, ProteinTech 66008-3-Ig) and monoclonal mouse anti-HA antibody (1:500, Invitrogen 26183), respectively, in combination with goat anti-mouse-HRP conjugate IgG (1:2000, ProteinTech SA00001-1). Blots were developed by addition of HRP substrate (Pierce ECL Western Blotting Substrate) and visualized with the Bio-Rad Gel Doc XR+ System (upper right panel). Blots were quantified by ImageJ 1.53e and band signal intensities were standardized to the Coomassie-signal of the complete lane. Relative abundance was quantified as the ratio of the standardized signal intensity at 60 min of 86°C to the standardized signal intensity 75°C (bottom right panel). **b** Spot test to validate survival. Culture samples at 75 °C and 60 min of heat shock were diluted to $OD_{600nm}$ 0.1 (= $10^{-1}$ dilution) based on the $OD_{600nm}$ measurement at the start of the experiment. Ten µL of a dilution series ($10^{-1}$ – $10^{-6}$) was spotted on gelrite plates. Plates were left to incubate for 5 days at 75°C for the regeneration of heat shock surviving cells. One representative 75°C-sample is shown.

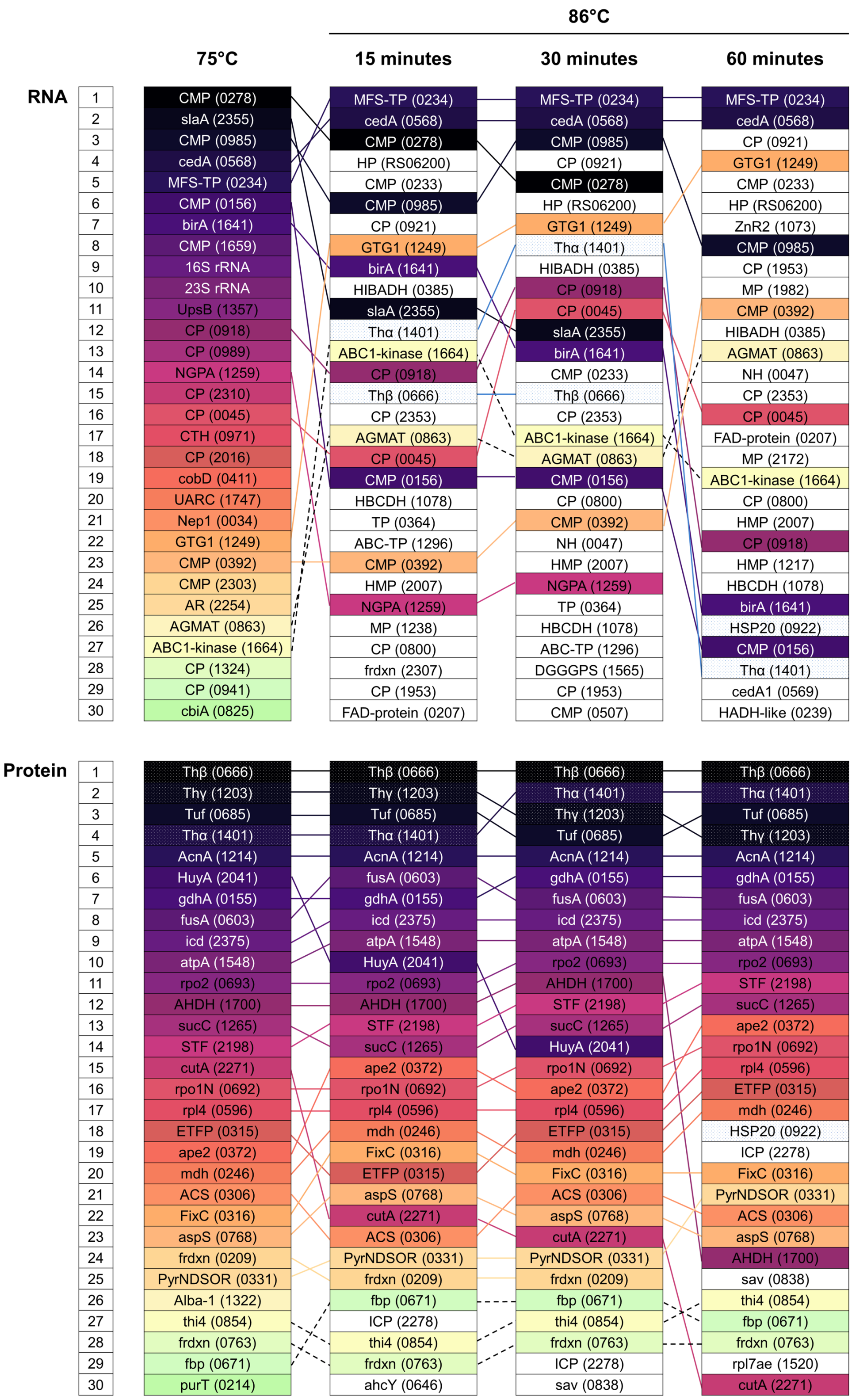


**Supplementary Figure S4. Top 30 of the most abundant transcripts and proteins in exponentially growing *S. acidocaldarius* at 75 °C and upon heat shock at 86 °C as determined by RNA-seq and MS.** Transcripts and proteins are ranked according to decreasing normalized CPM value or counts, respectively, and are colored according to their rank at 75 °C. Locus tags (Saci_xxxx) are indicated in between brackets. CP = conserved protein. CMP = conserved membrane protein. MP = membrane protein. HP = hypothetical protein. TP = transporter. The most abundant RNA species in the cell corresponds to the rRNA. However, due to the rRNA-depletion step in the RNA-seq workflow, counts for the rRNA species are low.

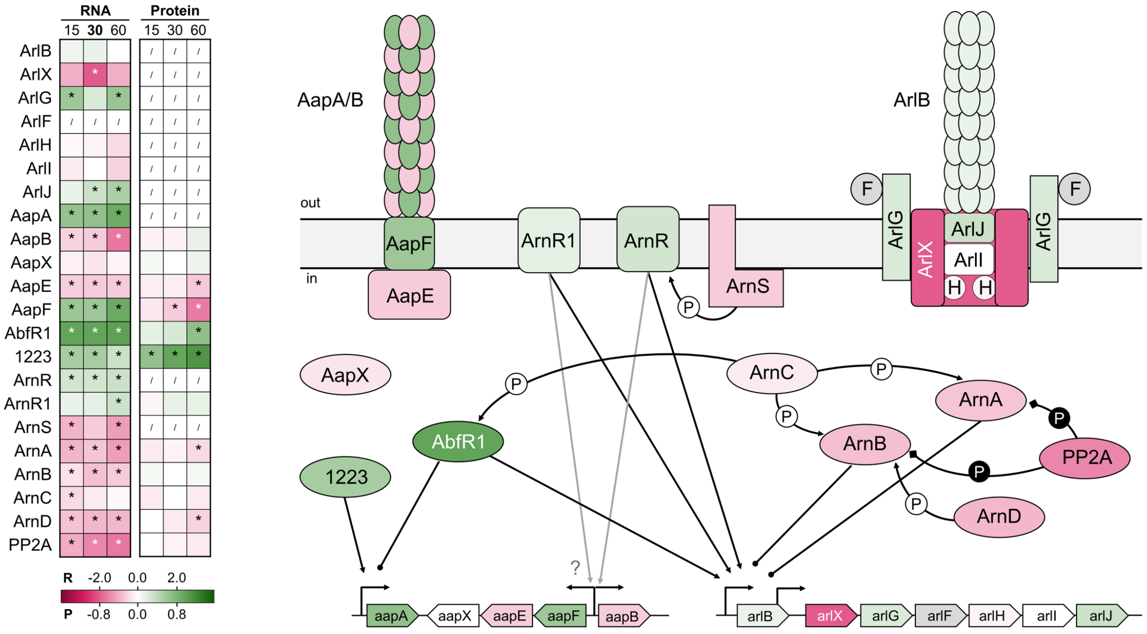


**Supplementary Figure S5. Heat-shock responsive differential expression of the aap pilus and archaellum.** The inset table shows differential expression at the RNA (R) and protein (P) level at all time points (15, 30, 60 minutes), color-coded according to the $log_2FC$ value in the gradient below. * = significant (FDR/adj.p-value < 0.05). / = not covered. The aap pilus is involved in biofilm formation and consists of AapA and AapB pilins, anchored in the membrane by the AapF and AapE motor proteins (29). AapX, a putative iron-sulfur oxidoreductase, is encoded in the same gene cluster (29). The archaellum is the motility structure, of which all seven structural components of the archaellum (ArlB, ArlX, ArlF-J) are encoded in an operon and of which transcription is controlled by two promotors, upstream of *arlB* and upstream of *arlX* (72). Regulation of both cellular processes is intertwined. The Lrs14-type AbfR1 is the major archaellum transcriptional activator and biofilm repressor (31). The membrane-bound one-component transcription factors ArnR and its paralog ArnR1 are activators for archaellum transcriptional activation upon, e.g. nutrient starvation (73), and are involved in regulating aap-expression. The transcriptional regulators ArnA and ArnB negatively regulate archaellum-transcription and are phosphorylated and activated *in vivo* by the kinases ArnC (phosphorylating ArnA and ArnB) and ArnD (phosphorylating ArnB only) and can be dephosphorylated by the PP2A Ser/Thr phosphatase (54, 71). The scheme on the right is colored according to differential expression at the RNA level at 30 minutes and is adapted from (19, 22). Arrow = positive regulation (activation), dot = negative regulation (repression). P on white background = phosphorylation, P on black background = dephosporylation.

**a**

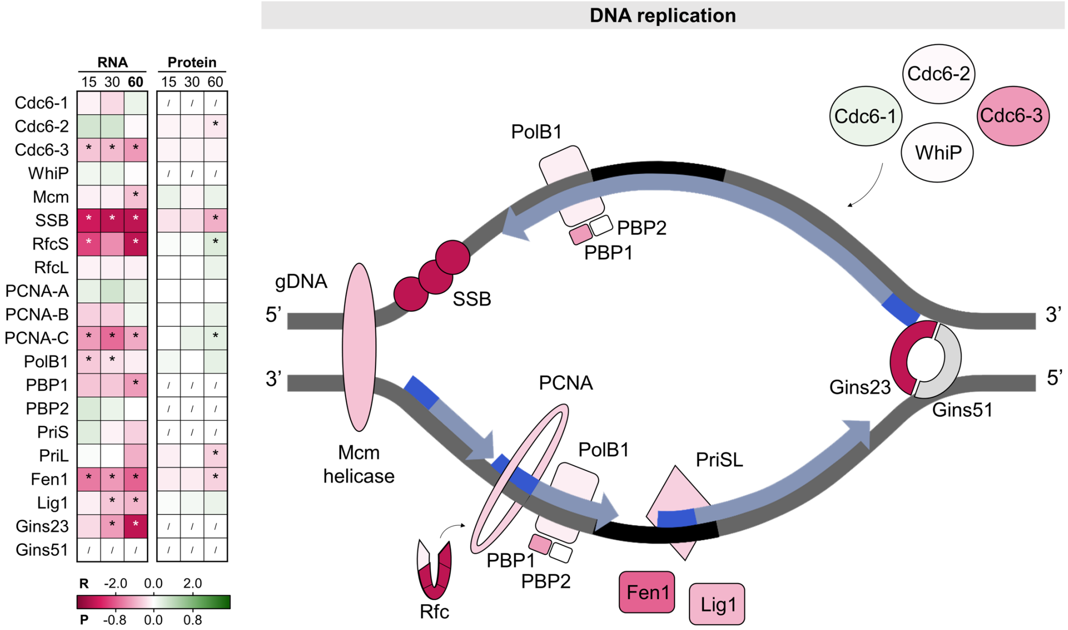


**b**

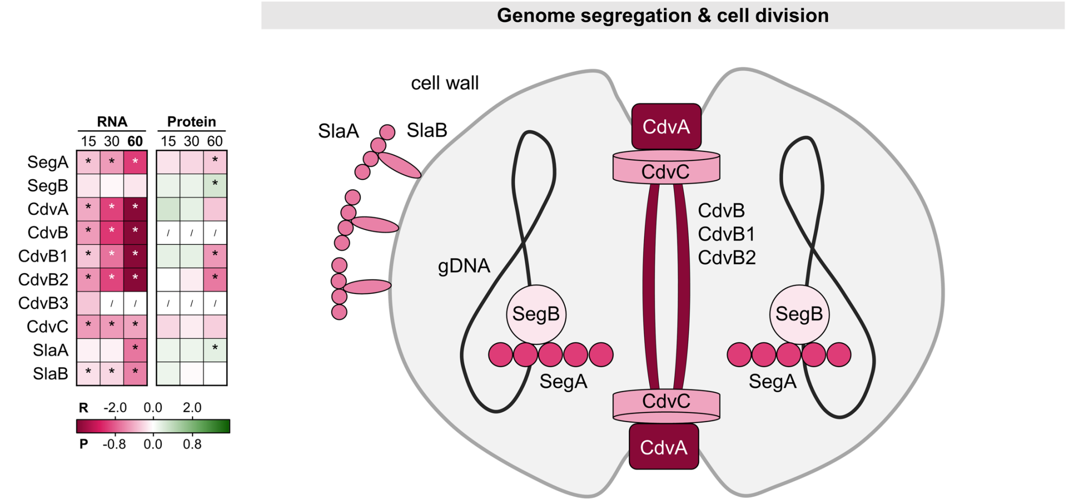


**Supplementary Figure S6. Heat-shock responsive differential expression of DNA replication, genome segregation and cell division machineries.** The table shows the DE of the genes involved at the RNA (R) and protein (P) level at all time points after heat shock (15, 30, 60 minutes), color coded according to the $log_2FC$ value in the gradient below. * = significant (FDR/adj.p-value < 0.05). / = not covered. The figure shows the proteins in their cellular context, colored according to DE at the RNA level after 60 minutes. **a** Differential expression of DNA replisome upon heat shock. DNA replication is initiated by binding of a replication initiator protein (Cdc6-1, Cdc6-2, Cdc6-3 and WhiP) to its corresponding origin(s) (35), followed by recruitment of the replisome and Mcm DNA helicase, which unwinds the DNA helix and forms a single-stranded (ss) DNA template (38). The single-strand DNA binding protein (SSB) is involved in binding and thereby protecting single-stranded DNA which is exposed during a variety of DNA-based processes (27, 32) The trimeric ring-shaped proliferating cell nuclear antigen (PCNA) (75) is loaded onto the DNA by the replication factor C (Rfc) complex and attaches the main replicative DNA polymerase PolB1 to the DNA template (27, 34, 73). For the lagging DNA strand, the DNA primase PriSL generates RNA primers, which are removed by the nuclease Fen1, followed by Okazaki fragment ligation by DNA ligase Lig1. Additional support of the replisome is provided by the Gins complex (Gins23-Gins51)(38). **b** Differential expression of the genome segregation and cell division machineries upon heat shock. Genome segregation is mediated by SegA and SegB, with SegB binding the chromosome and SegA polymerization separating the two DNA copies (27; 28) Cell division is initiated by CdvA ring-like positioning at the midcell membrane, which triggers formation of a non-contractile CdvB ring at the future division site, the latter forming a scaffold for the assembly of the CdvB1/B2 ring. Proteasomal degradation of CdvB allows for constriction of the CdvB1/B2 ring and cytokinesis (41, 44, 70, 80). Accessory roles are played by CdvB3 and CdvC (40). The cell wall of *Sulfolobales*, the S-layer, is composed of SlaA, anchored in the cytoplasmic membrane by SlaB (67). The scheme on the right is colored according to differential expression at the RNA level at 60 minutes and is adapted from (41, 70).

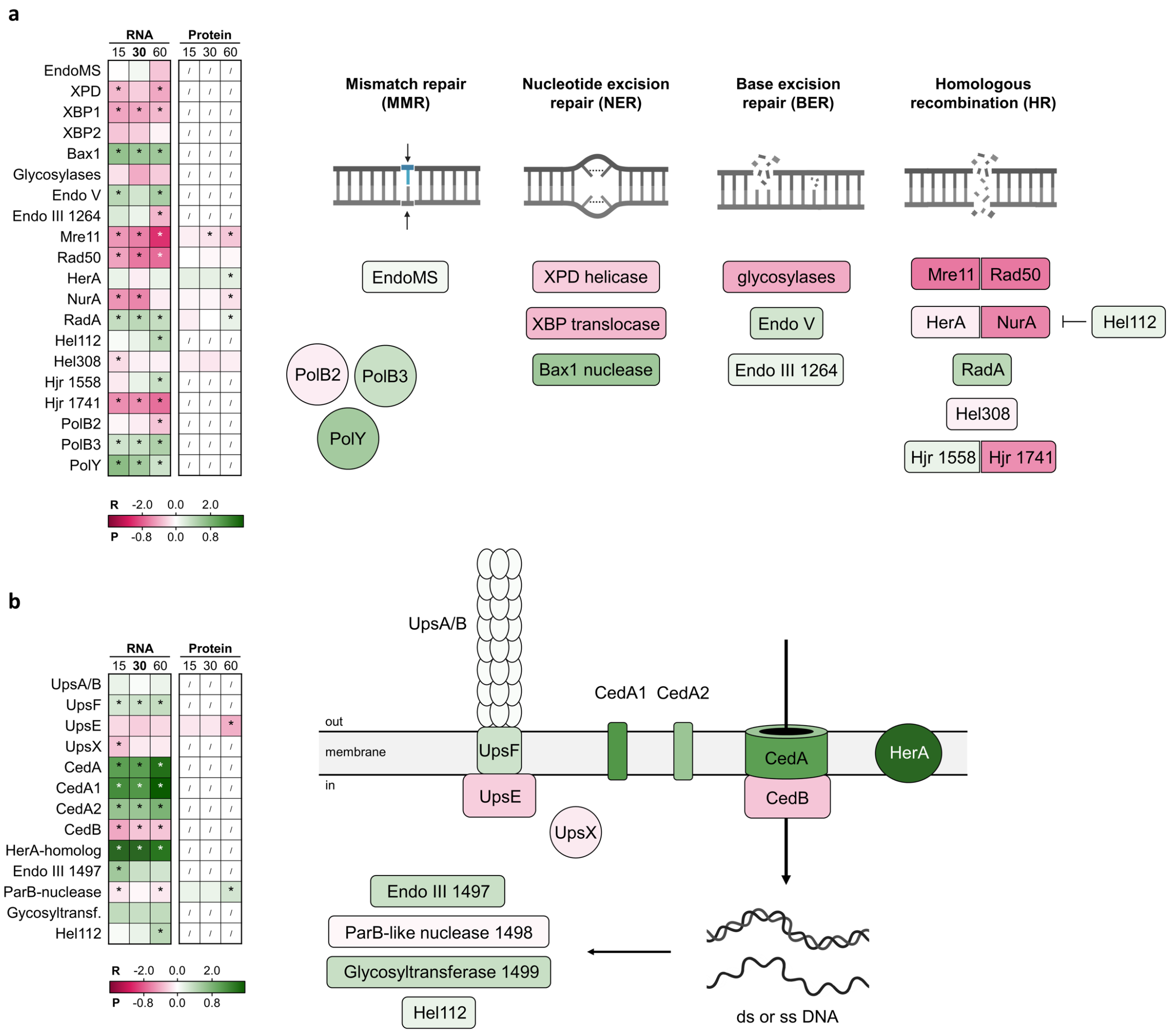


**Supplementary Figure S7. Heat-shock responsive differential expression of the DNA repair mechanisms and the Ced DNA import system.** The table shows the DE of the genes involved at the RNA (R) and protein (P) level at all time points after heat shock (15, 30, 60 minutes), color coded according to the $log_2FC$ value in the gradient below. * = significant (FDR/adj.p-value < 0.05). / = not covered. The figure shows the proteins in their cellular context, colored according to DE at the RNA level after 30 minutes. **a** Differential expression of the universally conserved DNA repair pathways. The three accessory DNA polymerases (PolB2, PolB3 and PolY) are involved in DNA repair and DNA damage tolerance (47, 48). Direct lesions of the DNA are mostly repaired by mismatch repair (MMR), nucleotide excision repair (NER) or base excision repair (BER), in which the affected nucleotides are removed and resynthesized using the undamaged DNA strand as template (47). Mismatched nucleotides generated during DNA replication are detected and repaired by EndoMS in MMR (50, 52). NER is involved in the repair of helix-destabilizing lesions induced by photoproducts. The XPD helicase is essential for DNA unwinding, and the complex consisting of XPB dsDNA translocase and Bax1 endonuclease assists in repair (50, 81). *S. acidocaldarius* encodes two XPB paralogs (XPB-1 and XBP-2). Hydrolytic depurination of the DNA, deamination of cytosine, oxidation or methylation is corrected by BER. In the canonical BER pathway, a glycosylase specific for the damage base detects the lesion and cleaves the glycosidic bond (51) and AP endonuclease Endo III (52) detects an abasic nucleotide and cleaves the DNA backbone, allowing a DNA polymerase to initiate DNA repair synthesis and DNA ligase to ligate the nick. Additional processing might occur by flap displacement and excision by Fen1 nuclease (50, 53). PCNA may also be involved in this process (53). In the alternative BER pathway, endonucleases such as endoV are involved in nicking the DNA next to the DNA lesion instead of a glycosylase (47). In Sulfolobus, repair of DNA double stand breaks (DSBs) is mainly established by HR and requires the presence of a second intact DNA copy (47). Repair is initiated by DSB end recession by the 5’ to 3’ exonuclease complex Mre11-Rad50 and helicase/nuclease complex HerA-NurA, yielding 3’ ssDNA ends which form nucleoprotein filaments with RadA recombinase, after displacement of SBB (47). The RecQ-like helicase Hel112 inhibits NurA-HerA nuclease activity (79). The nucleoprotein filament engages in an homology search with the intact DNA copy and catalyzes strand exchange and the formatting of a displacement (D-) loop. The 3’end of the invading template strand in the D-loop is either unwound by the helicase Hel308 or the D-loop may capture the second end of the DNA DSB, forming a four-way Holliday junction, which is then resolved by Holliday junction resolvases (Hjr) and possibly inducing cross-over (47). **b** Differential expression of the Ups-pilus and the Ced

DNA import system upon heat shock. The ups pili, built from UpsA and UspB pilin subunits and anchored in the membrane by UspF and ATPase UpsE (59), are involved in species-specific cellular aggregation (58). The Ced-system consists of four transmembrane proteins (CedA, CedA1, CedA2, CedB) and a putative Ced-associated ATPase (HerA-homolog). CedA and CedB are required for chromosomal DNA import, either in a ssDNA or dsDNA form (60). An ups-neighboring gene cluster, consisting of the Endo III, a ParB-like nucleases, a glycosyltransferase and helicase Hel112, is involved in subsequent DNA processing (54). Note that it is still unknown how DNA export is established (60). Scheme adapted from (62, 63).

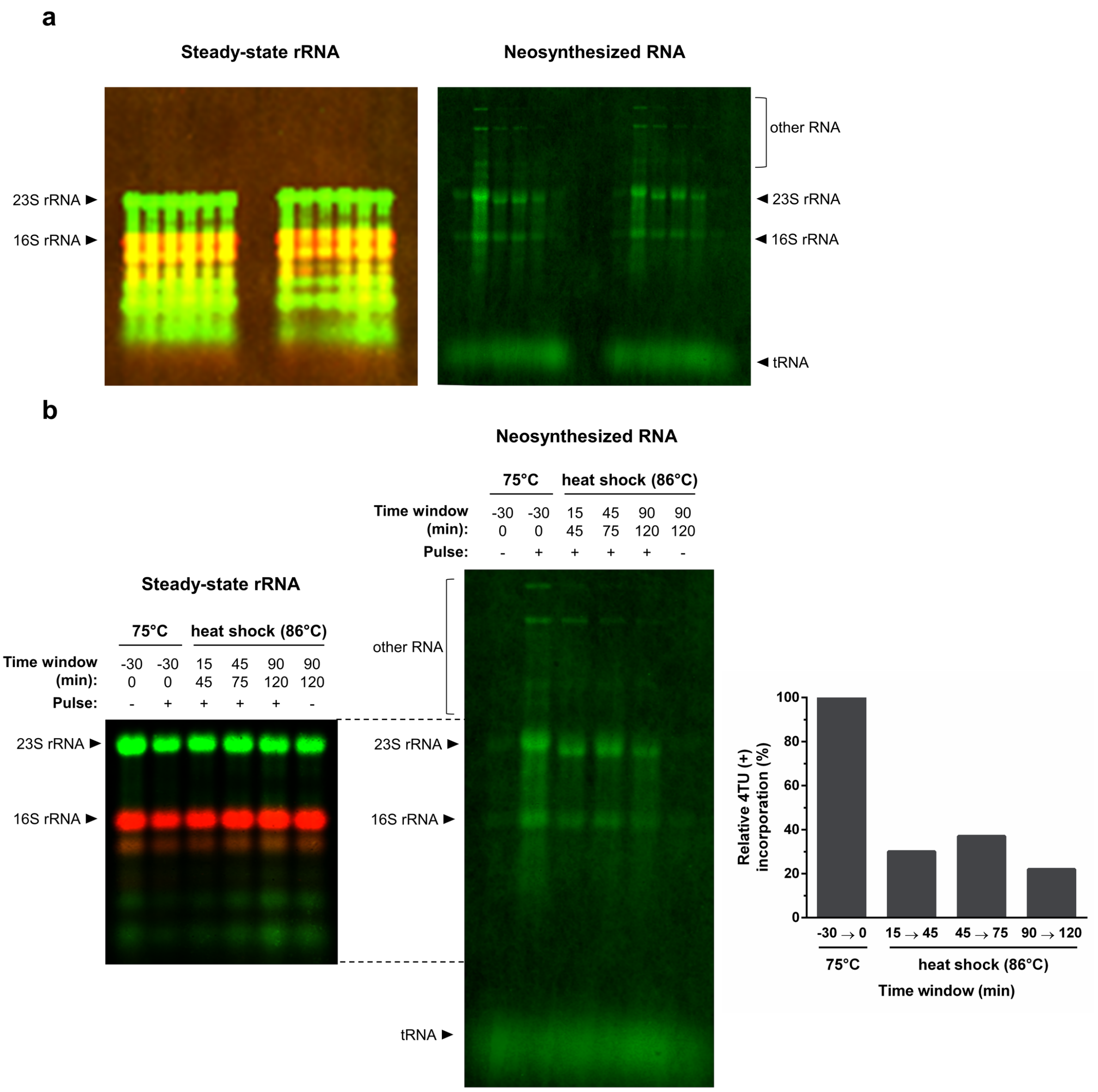


**Supplementary Figure S8. Pulse-labeling of neosynthesized RNA in exponentially growing *S. acidocaldarius* at 75°C and upon heat shock, extended set of data. a** Scan of the complete membrane of all samples. The large RNA species which incorporated the 4TU-label, detected by DyLight800-conjugated Streptavidin (right) do not have a rRNA origin since they are not recognized by the DY682- or DY782-coupled probes targeting the 5'end region of mature 16S or 23S rRNA (overexposed scan, left). **b** Pulse-labeling of neosynthesized RNA, extended set of heat shock time windows until 120 minutes *post* heat shock. SK-1 Thα-FLAG + Thβ-6xHis + Thγ-HA cultures were pulsed, both at 75°C and upon a 86°C heat shock for 30-minute time windows by addition of an excess of nucleotide analog 4TU (+) or uracil (-) as mock-control. 4TU was incorporated into neosynthesized RNA and 4TU-labeled RNA was biotinylated in the total pool of RNA. Total RNA was analyzed by Northern blotting. Bulk steady-state rRNA was detected by DY682- or DY782- coupled probes targeting the 5'end region of mature 16S or 23S rRNA, serving as a loading control (left panel). 4TU-labeled RNA was detected by DyLight800-conjugated streptavidin (central panel). Incorporation of 4TU into 16S or 23S rRNA was quantified by first subtracting the background-signal in the uracil-mock control and dividing this by the signal of the steady-state rRNA. Relative 4TU-incorporations upon heat shock were calculated relative to incorporation at 75°C and the average of 16S and 23S rRNA was determined as a ratio to express the relative transcriptional activity upon heat shock (right panel).

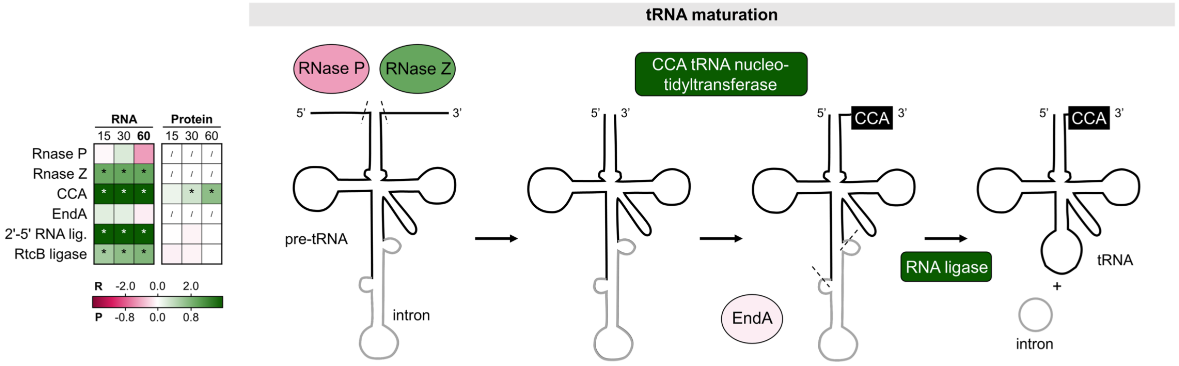


**Supplementary Figure S9. Heat-shock responsive differential expression of genes involved in tRNA maturation.** The inset table shows differential expression at the RNA (R) and protein (P) level at all time points (15, 30, 60 minutes), color coded according to the $log_2FC$ value in the gradient below. * = significant (FDR/adj.p-value < 0.05). / = not covered. The scheme on the right is colored according to differential expression at the RNA level at 60 minutes. Generation of mature tRNA requires many processing steps, initiated by removal of 5' leader sequence by RNase P and 3' trailer sequence by RNase Z from the pre-tRNA (80). Then, the CCA tRNA nucleotidyltransferase builds and repairs the 3'-terminal CCA sequence of tRNAs (83, 84). About half of Crenarchaeal tRNA genes contain introns, which are removed by a RNA splicing endonuclease EndA, followed by ligation of the splice products by a RNA ligase and finally yielding a mature tRNA (83, 85).

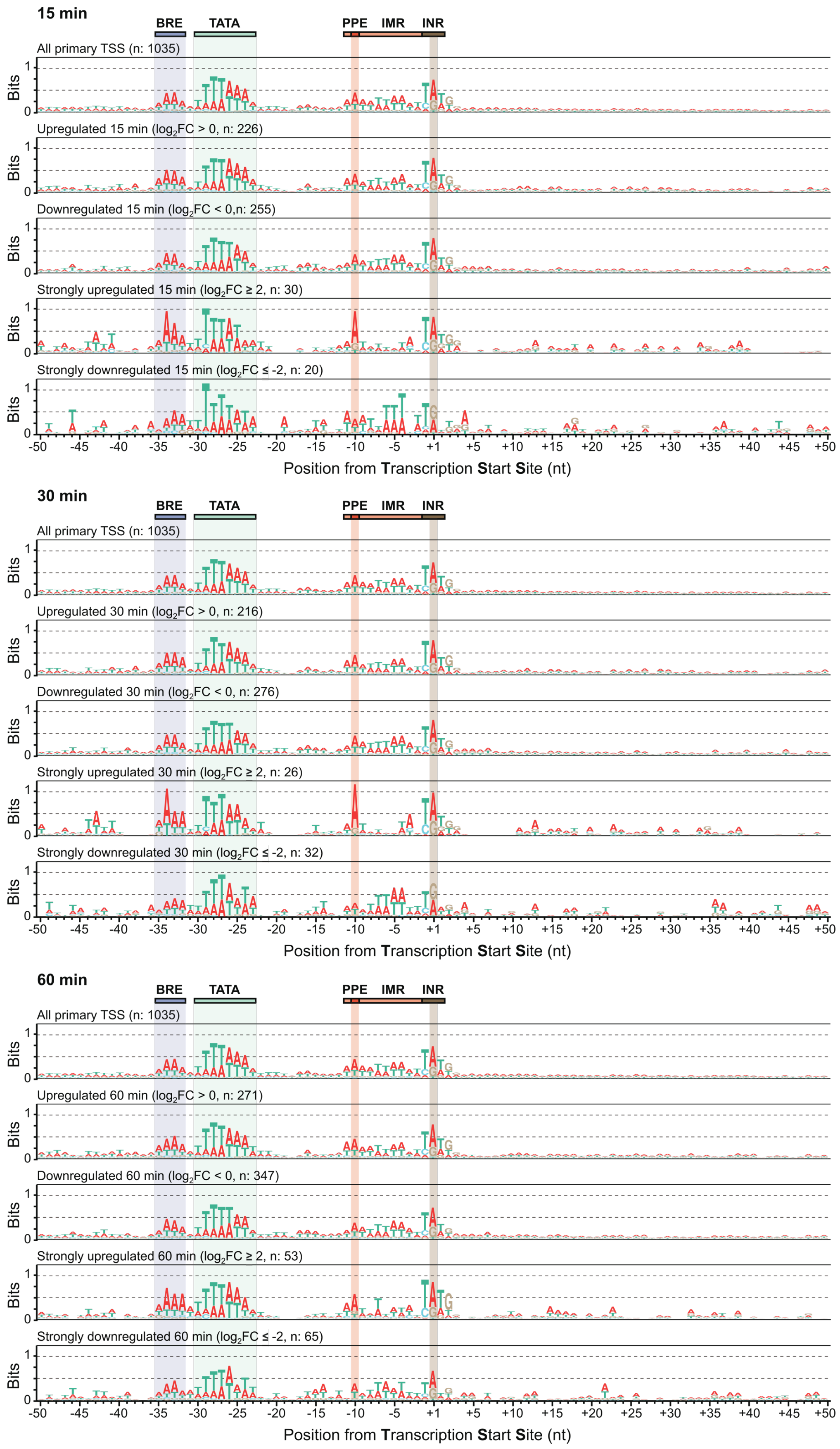


**Supplementary Figure S10. Extended motif of the promoter and 5' end of primary transcripts and subsets of regulation groups at 15, 30 and 60 minutes after heat shock.** PPE = proximal promoter element. IMR = Initially melted region. INR =Initiator element.

#### Supplementary Datasets (separate Excel-files)

**Supplementary Dataset S1.** Overview of cultures and samples in omic experiments.

**Supplementary Dataset S2.** RNA-seq processing parameters and sequencing stats.

**Supplementary Dataset S3.** Overview of RNA-seq and MS data.
